# Translational repression of viral RNA mediates arbovirus persistence in mosquitoes

**DOI:** 10.1101/2025.07.10.664053

**Authors:** Marc Talló-Parra, Mireia Puig-Torrents, Gemma Pérez-Vilaró, Sol Ribó Pons, Xavier Hernández-Alias, Federica Mantica, Manuel Irimia, Luis Serrano, Juana Díez

**Author notes:** Co-first authorship (Marc Talló-Parra, Mireia Puig-Torrents). Corresponding autor.

## Abstract

Persistent infection of mosquito cells is essential for the transmission of arboviruses, yet how these viruses produce sufficient progeny without compromising host cell fitness remains unclear. Arbovirus genomes exhibit suboptimal codon usage for both human and mosquito hosts, raising questions about how they achieve efficient translation in such distinct cellular environments. Using chikungunya virus (CHIKV) as a model, we conducted a temporal, genome-wide analysis of transcription, translation, and tRNA modifications in *Aedes albopictus* C6/36 cells. Unlike in human cells, CHIKV infection does not alter the tRNA modification landscape in mosquitoes to overcome codon bias, nor does it induce widespread degradation of host transcripts. Instead, viral persistence is marked by a progressive, virus-specific repression of CHIKV RNA translation, occurring alongside a recovery of host mRNA translation. This translational balance, maintained independently of the RNAi system, enables sustained viral production without major disruption to host gene expression. Our findings identify translational control as a central mechanism underlying persistent arbovirus infection in mosquito cells.

## Introduction

Arboviruses transmitted by mosquitoes, including dengue (DENV), West Nile (WNV), Zika (ZIKV) and chikungunya (CHIKV) viruses, represent major global health threats [1]. In recent decades, urbanization, climate change and globalization have facilitated the spread of these viruses and their mosquito vectors beyond tropical and subtropical regions, reaching new geographical areas worldwide. Today, more than 80% of the global population is at risk of infection by at least one of these viruses, with outbreaks increasingly reported in European countries [2]. Despite the urgent need for effective control measures, vaccines and antiviral therapies remain unavailable for most mosquito-borne viruses [3]. As a result, limiting viral transmission through mosquito control has become a central strategy for curbing the spread of these infections. However, traditional approaches, such as insecticide use and breeding site elimination, have proven insufficient to halt arbovirus spread. Although novel interventions like the release of *Wolbachia*-infected mosquitoes offer promise, their efficacy varies across ecological and geographical context [4]. Consequently, there is an urgent need for complementary innovative vector control strategies.

Mosquito-borne viruses exhibit a remarkable ability to propagate efficiently in both human and mosquito hosts, two organisms separated by over one billion years of evolutionary divergence. In humans, these viruses cause acute and often severe infections, whereas in mosquitoes, they establish life-long, persistent infections without inducing apparent pathological effects [5]. Because persistent infection in mosquitoes is essential for arbovirus transmission and propagation, understanding the mechanisms that support it is critical for developing novel vector control strategies. Yet, how arboviruses replicate successfully in such evolutionary distinct hosts, while producing drastically different outcomes, remains unsolved. Moreover, interactions between arboviruses and their mosquito hosts are far less characterized than those in vertebrates. A key step in the arbovirus replication cycle is the efficient expression of viral proteins, which relies entirely on the host cell machinery [6]. Interestingly, the codon usage of diverse arboviruses, including CHIKV, is suboptimal for both human and mosquito hosts. In both vertebrates and mosquitoes, highly expressed genes preferentially use G/C-ending codons, whose cognate tRNAs are highly abundant, whereas arbovirus genomes are enriched in A/U-ending codons, whose cognate tRNAs are scarce [7]. This codon bias is predicted to slow down translation elongation and decrease protein expression. However, using CHIKV as a model, recent studies show that in human cells, CHIKV infection overcomes this limitation by inducing alterations in the tRNA modifications landscape. Upon infection, CHIKV triggers a DNA damage stress response that increases the levels of the mcm^5^ tRNA modification. This change reprograms codon optimality, selectively enhancing the translation of specific suboptimal codons that are highly enriched in both host DNA damage stress response genes and CHIKV genomes. Hence, CHIKV codon usage optimally aligns with the tRNA modification landscape in human infected cells [8]. Whether a similar strategy operates in mosquito cells, and how these cells sustain viral protein synthesis without compromising their own viability remains unknown. Here, using CHIKV as a model, we address these fundamental questions by performing a temporal analysis of the transcriptome, translatome and tRNA modification landscapes in mosquito cells. In contrast to human cells, we found that CHIKV infection in mosquito cells does not remodel the tRNA modification landscape to promote viral protein expression. Instead, CHIKV persistence is characterized by a virus-specific translational repression that enables sustained, yet controlled, viral production without major disturbances to host mRNA translation. This translational repression is mechanistically linked to two key factors limiting CHIKV protein synthesis in mosquito cells compared to human cells: a codon optimality constraint and divergent functions of viral non-structural protein 2 (NSP2). Our results show a fine-tuned translational equilibrium in mosquito cells that allows both the sustained viability of mosquito cells and the establishment of CHIKV persistence through controlled viral protein synthesis.

## Results

### An imbalance between viral RNA and protein expression characterizes CHIKV persistence in mosquito cells

CHIKV is a positive-sense single stranded RNA virus whose genome structurally mimics host mRNAs, bearing a 5’ cap and a 3’ poly(A) tail. The CHIKV RNA genome (gRNA) contains two open reading frames (ORFs). The first ORF is translated directly from the genome and encodes the non-structural proteins (NSPs) required for viral replication, while the second ORF is expressed from a subgenomic RNA (sgRNA) transcribed during replication and encodes the structural proteins (SPs) necessary for virion assembly (**Fig 1A**). To investigate how CHIKV establishes persistent infection in mosquito cells without major effects on host cell physiology, we first analyzed CHIKV infection kinetics in *Aedes albopictus* C6/36 cells to identify the time point at which viral replication plateaus, referred to hereafter as the time of persistence. We selected the C6/36 cell line, which lacks a functional RNA interference (RNAi) system, to specifically investigate potential translational control mechanisms involved in viral persistence. With this, we avoid possible cross-effects derived from the RNAi system that could confound the interpretation of our results. Cells were infected at a multiplicity of infection (MOI) of 4, a condition under which nearly all cells become infected after a single replication cycle (**S1 Fig**). Samples were collected at early (3, 6, 8 hours post-infection (hpi)) and late time points (1, 2, 5 and 7 days post-infection (dpi)). Viral titers (**Fig 1B**) as well as gRNA and sgRNA levels, together with their encoded proteins (**Fig 1C-D**), were assessed. Interestingly, viral titers and protein levels exhibited distinct kinetics compared to gRNA and sgRNA levels. Viral titers and protein expression peaked around 1-2 dpi and subsequently declined. In contrast, gRNA levels continued to rise, reaching a maximum at 5 dpi while sgRNA plateaued as early as 8 hpi and remained stable until 5 dpi, after which a slight decrease was observed, as for gRNA. The decline in viral titers and protein levels despite sustained high viral RNA levels suggests the presence of post-transcriptional regulatory mechanisms, either translational regulation or sequestration of gRNA and sgRNAs in compartments inaccessible to RNAses and to the translation machinery. To determine whether this observation was influenced by the absence of a functional RNAi system, we analyzed CHIKV replication kinetics in U4.4 cells, an *A. albopictus* cell line with a competent RNAi response. As expected, viral titers, RNA, and protein levels were lower in U4.4 cells than in C6/36 cells, consistent with RNAi-mediated antiviral activity **(S2 Fig**). However, the overall kinetics were comparable in both cell lines, indicating that similar regulatory mechanisms were operating in both, and that the observed uncoupling of viral RNA and protein levels is independent of the RNAi pathway. Therefore, all subsequent experiments were conducted using C6/36 cells.

**Fig 1.**
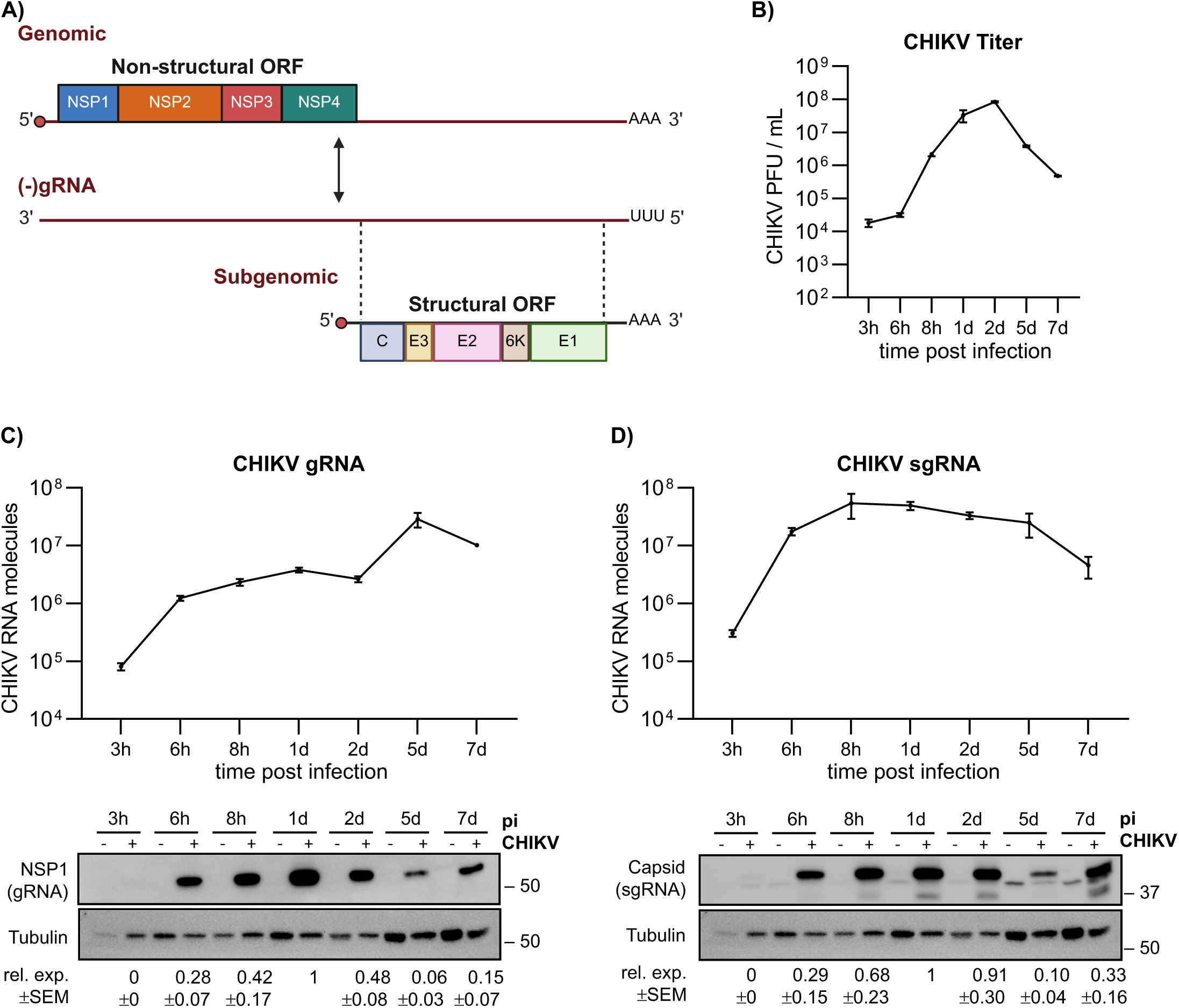
Kinetic analysis of CHIKV infection in C6/36 mosquito cells. (A) Scheme of CHIKV RNA genome and its protein synthesis strategy. (B) CHIKV titers over time, determined by plaque assay. (C) Infection kinetic of CHIKV gRNA levels determined by qPCR and NSP1 protein levels assessed by western blotting. Tubulin was used as loading control. (D) Infection kinetic of CHIKV sgRNA levels and capsid protein expression, analyzed as in panel (C). All infections were performed at a MOI of 4 and samples were collected at the indicated time post-infection. Data points represent the mean ± SEM of 3 independent biological replicates. Viral RNA levels were quantified using a standard curve of an in vitro generated CHIKV RNA. Protein expression levels correspond to the intensity quantification of each band normalized to tubulin levels and to the 1 dpi value within each blot.

### Translation of CHIKV RNAs is repressed in persistently infected mosquito cells

To directly address whether the distinct CHIKV RNA and protein kinetics were linked to translation alterations, we performed polysome profiling analyses. This technique assesses translation effects by measuring the distribution of global mRNAs based on ribosomal occupancy, distinguishing between untranslated RNAs (free or ribosomal subunits), monosome-bound RNAs (associated with a single ribosome), and polysome-bound RNAs (actively translated by multiple ribosomes). Based on CHIKV protein expression kinetics in C6/36 cells (**Fig 1 C-D**), we selected six time points: 3, 6, and 8 hpi, corresponding to the phase of increasing viral RNA and protein levels; 1 dpi, representing peak of protein expression; and 5 and 7 dpi, when viral protein levels decline despite sustained RNA abundance. Polysome profiles revealed that CHIKV infection induces a transient global translational repression, evidenced by an increased monosome fraction (around fractions 10–12) and a reduced polysome fraction (around fractions 13–20) between 3 and 8 hpi (**Fig 2A**). This repression gradually diminished over time, and by 7 dpi, the translational profiles of infected and control cells were indistinguishable, indicating recovery of basal translation. To quantify this effect, we calculated the monosome-to-polysome (M/P) ratio at each time point relative to non-infected controls (**Fig 2B**). Values >1 indicate higher levels of monosome than polysomes, indicating inefficient translation, whereas values <1 indicate the opposite. The M/P ratio confirmed a strong repression at 8 hpi, which progressively resolved by 5 dpi. To determine whether this repression affected viral RNAs or was restricted to host mRNAs, we measured the distribution of CHIKV gRNA and sgRNA across polysome fractions by qPCR (**Fig 2C**). Interestingly, viral gRNA and sgRNA displayed an opposite trend compared to global mRNAs: their M/P ratios progressively decreased up to around 1 dpi, indicating translational activation, and increased at later time points, reflecting a shift towards translational repression. Of note, sgRNA translation was considerably activated earlier than gRNA, likely reflecting the higher demand for structural proteins. Collectively, these results indicate that CHIKV RNA translation is repressed during the persistent phase of infection, linking viral persistence to reduced translational efficiency.

**Fig 2.**
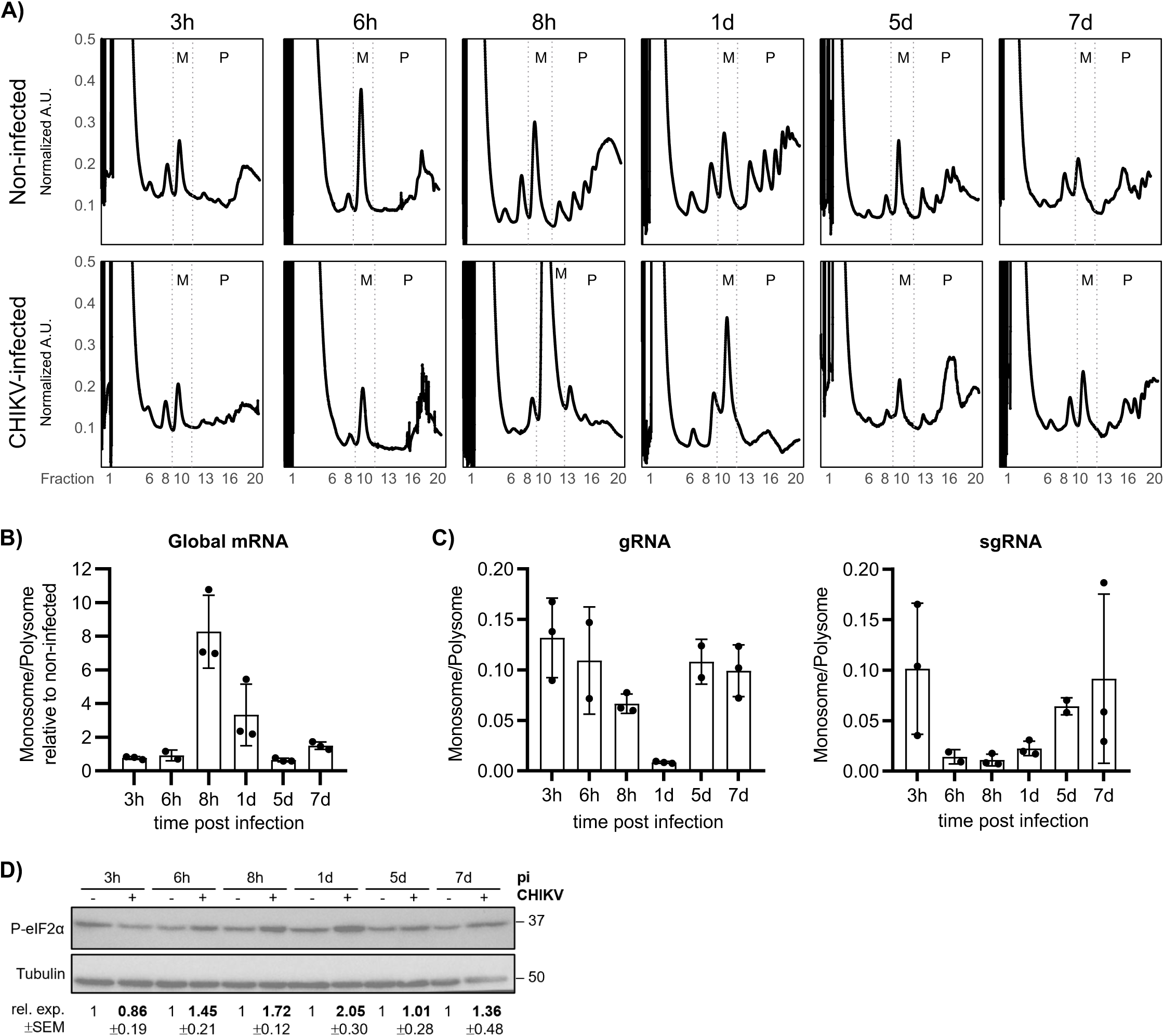
Dynamic translational changes during CHIKV infection. (A) Representative polysome profile analyses at different time points post-CHIKV infection. Profiles were normalized using BCA-based protein quantification. (B) Monosome to polysome ratios for global host mRNAs, calculated by measuring the area-under-the-curve (AUC) from polysome profiles and normalized to non-infected. (C) Monosome to polysome ratios for viral CHIKV gRNA and sgRNA, determined by qPCR of RNA extracted from polysome fractions. (D) Phospho-eIF2α protein levels during infection, assessed by western blotting. Tubulin was used as loading control. All infections were performed at a MOI of 4 and samples were collected at the indicated time post-infection. Bars represent the mean ± SEM of 3 independent biological replicates. Protein expression levels were normalized to tubulin and to the non-infected value within each blot.

To investigate the opposite translational behavior of host mRNAs and viral RNAs, we examined the phosphorylation status of eukaryotic translation initiation factor 2 alpha (eIF2α), a central event in the integrated stress response (ISR). Under stress conditions, including viral infection, eIF2α is typically phosphorylated, leading to an attenuation of translation initiation and global repression of protein synthesis [9]. However, many viruses interact with the ISR to either evade or manipulate it for their benefit in human and mosquito [10–12]. Whether CHIKV induces eIF2α phosphorylation in mosquito cells, and whether CHIKV RNAs can evade eIF2α-mediated translational repression remain unclear. To address this, we analyzed eIF2α phosphorylation at different time points post-infection. A transient increase in eIF2α phosphorylation was observed between 6 hpi and 1 dpi, coinciding with global host mRNA translation repression and viral RNA translation activation (**Fig 2D**). This indicates that CHIKV RNAs are capable of bypassing eIF2α-mediated translational inhibition. As the infection progressed into a persistent phase, eIF2α phosphorylation levels declined, host translation was restored, and CHIKV RNA translation became repressed.

### CHIKV infection alters transcriptional and translational landscapes of mosquito cells

To further investigate how CHIKV infection influences mosquito gene expression over time, we generated RNA-seq and Ribo-seq libraries to profile the host transcriptional and translational landscapes. As before, time points were selected to capture early stages of CHIKV infection (3, 6, 8, hpi), the peak of protein expression (1 dpi), and the persistent stage (7 dpi). Reproducibility between RNA-seq and Ribo-seq replicates was confirmed (**S3 Fig**) and quality metrics validated (**S4–11 Fig**). Across all samples, ribosome-protected fragments (RPFs) were within the expected size range (28–33 nt), exhibited clear triplet periodicity, showed no enrichment in untranslated regions (UTRs), and displayed a strong preference for frame 0 in both host and viral mRNAs. After quality assessment, we analyzed changes in the transcriptome and translatome during CHIKV infection. mRNA-seq analysis (**Fig 3A** and **S1 Table**) revealed that, in contrast to the drastic decrease in mRNA levels observed in CHIKV-infected human cells [8], CHIKV-infected mosquito cells exhibited a balanced transcriptional response, with a comparable number of mRNAs upregulated and downregulated. In human cells, CHIKV infection induces a global reduction in mRNA levels through the nuclear translocation of CHIKV NSP2, which promotes degradation of Rpb1, a core subunit of RNA Polymerase II (RNAPII), ultimately leading to transcriptional shutoff [13]. To determine whether this mechanism operates differently in mosquito cells, we examined the subcellular localization of NSP2 and found that, in contrast to human cells, it remains confined to the cytoplasm (**Fig 3B**). This cytoplasmic retention prevents transcriptional shutoff, thereby enabling a finely tuned, CHIKV-induced host transcriptional response (**Fig 3A**).

**Fig 3.**
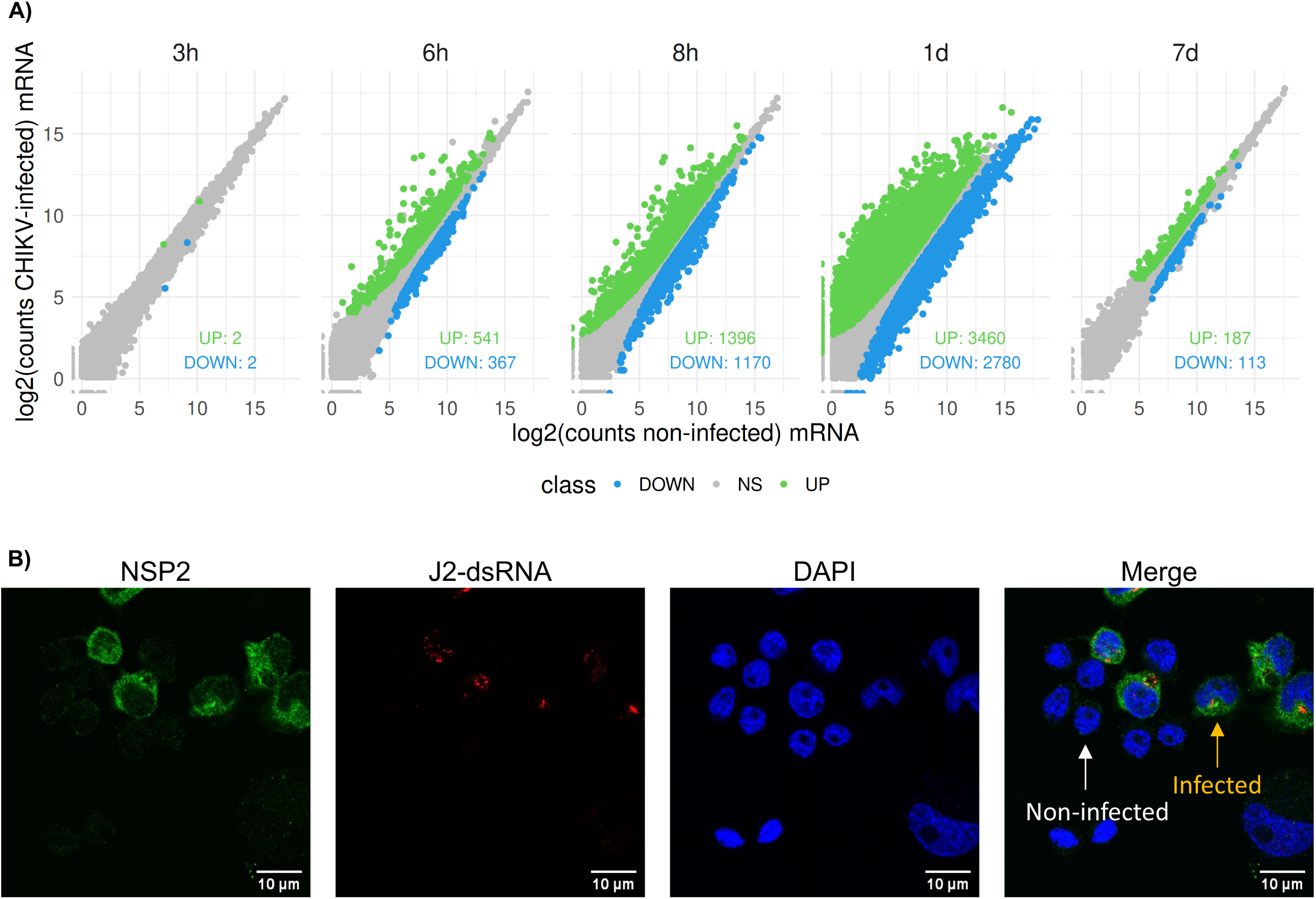
CHIKV infection alters dynamically the mosquito cell transcriptome. (A) Transcriptome analysis of host mRNAs in CHIKV-infected versus non-infected cells over time. Genes were plotted according to their log2 fold changes. Colors indicate genes that are significantly (p.adj < 0.05) upregulated (green) or downregulated (blue). (B) CHIKV-NSP2 (green) protein localization in C6/36 cells as determined by immunofluorescence. Infected cells were identified by labelling with J2 antibody (red), which recognizes dsRNA produced during CHIKV replication. Cells were stained with DAPI (blue) for visualization of nuclei. Scale bars correspond to 10 μm.

mRNA-seq analysis revealed a dynamic, time-dependent transcriptional response in CHIKV-infected mosquito cells. Minimal changes were detected at 3 hpi, however, the number of differentially expressed host mRNAs progressively increased over time, peaking at 1 dpi with approximately 6,000 transcripts either upregulated or downregulated. By 7 dpi, this number declined sharply, indicating a return toward baseline expression levels. Functional enrichment analyses (**S2 Table**) showed that during the early stages of infection (6 hpi to 8 hpi), upregulated mRNAs were enriched in gene-ontology (GO) terms associated with energy metabolism (glycolysis, amino acid metabolism, mitochondria), translation machinery (tRNA aminoacylation, ribosome components) and stress processes (unfolded protein binding, redox homoeostasis). Conversely, downregulated mRNAs were enriched in GO terms associated with DNA replication, repair, and nuclear organization, suggesting a halt in cell division and a shift of resources toward managing cellular stress. These changes are indicative of an intense stress that triggers the ISR. By 1 dpi, coinciding with the peak of virus production, the host transcriptional response shifted. While general metabolic activities remained upregulated, and DNA associated terms continued to be downregulated, a notable repression of transcripts related to mitochondrial oxidative phosphorylation, ribosome biogenesis, and RNA processing was observed, pointing to a metabolic transition from mitochondrial-dependent energy production to catabolic pathways. At 7 dpi, the number of differentially expressed transcripts decreased substantially, indicating a recovery from the initial ISR and a reactivation of most cellular functions. However, persistent upregulation of transcripts associated with glycolysis, the proteasome complex, and oxidoreductase activity points to maintenance of a residual oxidative stress. Collectively, these findings highlight a dynamic transcriptional response to CHIKV infection in mosquito cells, initially characterized by an acute stress response and cellular shutdown, followed by a metabolic reprogramming and a gradual progression towards homeostasis.

Next, we analyzed changes in the translatome by comparing ribosome-protected fragment (RPF) levels to mRNA abundance. First, we assessed the proportion of host and viral reads in both RNA-seq and Ribo-seq libraries. In the RNA-seq library, the proportion of viral reads steadily increased, reaching 63.62% of total reads at 1 dpi before declining to 16.12% by 7 dpi (**Fig 4A, top left**). In contrast, Ribo-seq analyses revealed that viral reads peaked earlier, at 32.92% by 8 hpi and dropped to 4.86% at 7 dpi (**Fig 4A, bottom left**). This discrepancy between viral RNA abundance and ribosome occupancy suggests translation repression of viral RNAs, consistent with results obtained from complementary techniques (**Fig 1** and **Fig 2**). To contextualize these findings, we compared them to previously published data from CHIKV-infected HEK293T human cells [8], where the cytoplasmic and endoplasmic reticulum-associated translatomes were analyzed separately. For direct comparison, we combined reads from both compartments (see Materials and Methods). In human cells, viral reads accounted for 28.14% of the RNA-seq library (**Fig 4A, top right**) and 58.55% of the Ribo-seq library (**Fig 4A, bottom right**), indicating a robust translational activation of viral RNAs. Taken together, these results demonstrate that in comparison to human cells, CHIKV RNAs in mosquito cells are not efficiently translated at any time point, even at the peak of viral RNA accumulation.

**Fig 4.**
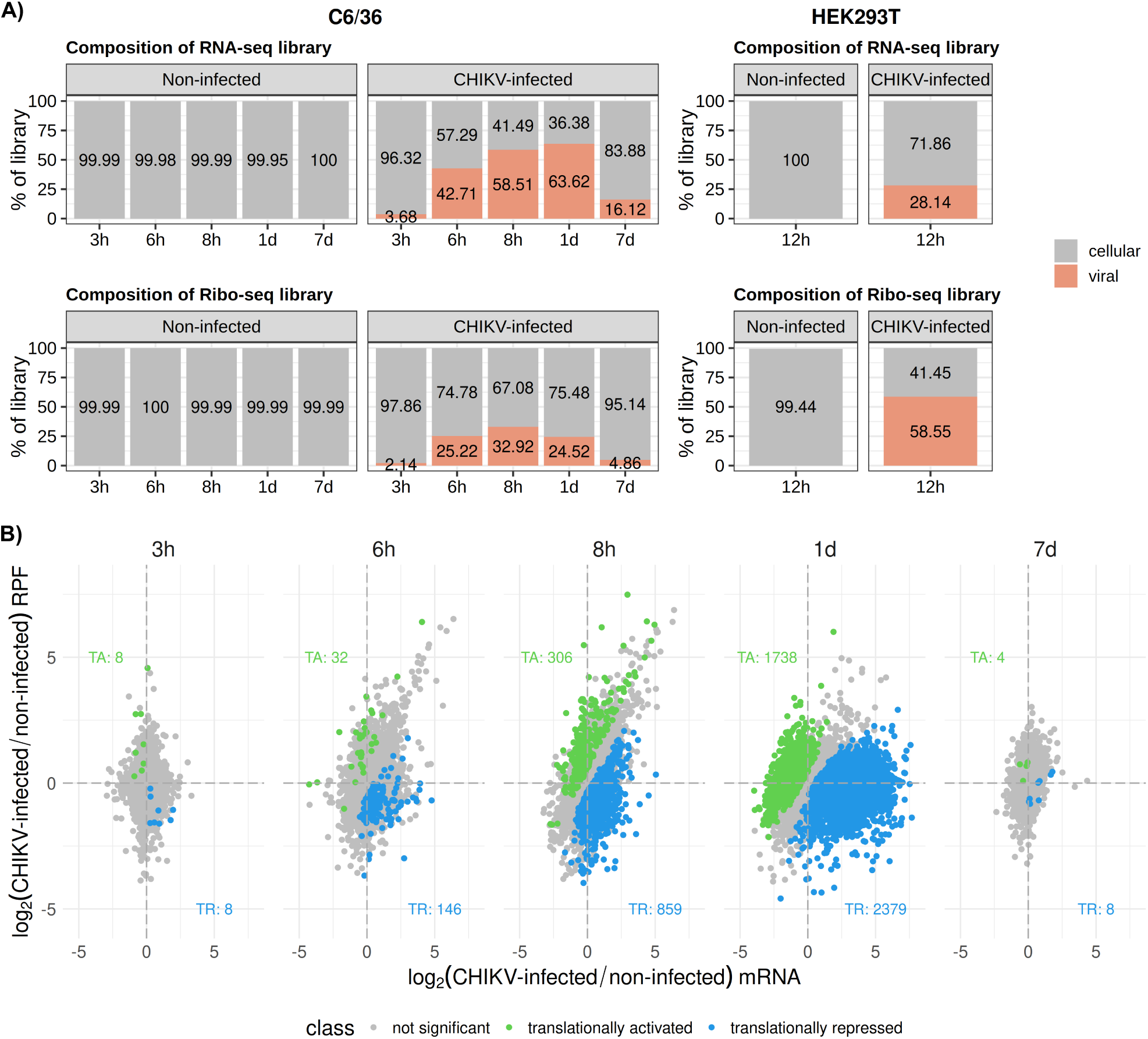
CHIKV infection dynamically alters the mosquito cell translatome. (A) Composition of RNA-seq and Ribo-seq libraries from non-infected and CHIKV-infected cells in C6/36 and HEK293T [8]. Displayed are the percentages of total cellular and viral raw counts. (B) Analyses of RNA-seq and RPF-seq libraries comparing CHIKV-infected and non-infected samples over time. Genes were plotted according to their log2 fold changes and colored based on significantly (p.adj < 0.05) changed translational efficiency: activated (green) or repressed (blue).

Next, translational efficiency (TE) was calculated for each host mRNA by dividing the normalized mean RPF counts by the normalized mean RNA-seq counts using RiboDiff [14]. Estimated significant changes represent alterations in RPF levels that could not be attributed to corresponding changes in mRNA abundance (**Fig 4B** and **S1 Table**). TE analyses revealed a dynamic, time-dependent remodeling of the host cell translatome. From 6 hpi onward, translational repression progressively intensifies. Similarly to transcriptional changes, the most significant shift in TE occurred at 1 dpi, and by 7 dpi the translatome largely recovered to baseline levels comparable to those observed in non-infected cells. Functional enrichment analysis (**S2 Table**) revealed that at 6 and 8 hpi, translationally repressed transcripts were enriched in categories related to translation (tRNA aminoacylation, ribosome components), metabolic pathways (glycolysis and amino acid metabolism) and protein folding. Notably, all these were transcriptionally upregulated indicating an uncoupling of the host transcriptional and translational responses, and a tight gene expression regulation aligned with eIF2α phosphorylation (**Fig 2D**). Additional enriched terms among translationally repressed transcripts include ATP activities and epigenetic regulators, both consistent with energy conservation and suppression of gene expression. By 1 dpi, a shift in translational control occurred. Translationally activated transcripts were primarily associated with translation machinery (ribosome components, eukaryotic preinitiation complex), mitochondrial function and redox homeostasis, transcription and RNA processing (DNA polymerase activity, splicing). These changes indicated that cells enter a phase of high metabolic activity, likely to support the increased demand for CHIKV protein synthesis. Concurrently, a selective translational repression of signaling pathways, regulatory mechanisms (e.g., phosphorylation, ubiquitin-mediated processes), and cytoskeletal/transport functions were detected. Interestingly, this translational shift contrasted with the transcriptional response. While transcription of genes involved in mitochondrial function, translation, and RNA processing was downregulated, their translation was selectively activated, suggesting a translation-driven enhancement of energy production to sustain cellular reorganization. By 7 dpi, the absence of major changes in gene expression indicate that the infection had reached a steady state, with global host translation largely restored. Overall, the combined transcriptomic and translatomic analyses reveal a complex, dynamic host response to CHIKV infection in mosquito cells. This response is characterized by an early ISR-mediated shutdown of non-essential functions, followed by a metabolic reprogramming phase, and ultimately a state of adaptation and recovery, despite evidence of oxidative imbalance.

The observed eIF2α phosphorylation pattern during CHIKV infection in mosquito cells, which peaked at 1 dpi (**Fig 2D**), indicated the activation of the ISR. While this key phosphorylation event broadly inhibits protein synthesis, it permits the preferential translation of Activating Transcription Factor 4 (ATF4). As a key transcriptional effector, ATF4 typically upregulates genes for stress adaptation and recovery, or for inducing cell death via activation of pro-apoptotic genes [15]. In agreement with activated ISR, the *A. albopictus* ATF4 ortholog was translationally activated from 6 hpi to 1 dpi, correlating precisely with eIF2α phosphorylation pattern, and returned to baseline levels by 7 dpi. Moreover, consistent with a pro-survival ATF4-mediated transcriptional response, downstream-ATF4 transcriptionally upregulated targets did not include pro-apoptotic genes, instead, anti-apoptotic genes BAG5 and BAG6 were transcriptionally upregulated, together with stress adaptation genes such as asparagine synthetase (ASNS) and tribbles pseudokinase 3 (Trib3), involved in metabolic adaptation, and genes involved in antioxidant defense, such as the master activator NRF2/Cnc and glutamate-cysteine ligase catalytic subunit (GCLC). Collectively, these results indicate that ATF4 activation during the initial stages of CHIKV infection in C6/36 cells primarily modulates metabolism and antioxidant activity, favoring cell survival.

Previous studies in human cells have shown that CHIKV infection activates a DNA damage-induced stress response, leading to elevated levels of the mcm⁵ tRNA modification. This shift alters codon optimality, enhancing the translation of suboptimal codons enriched in both the CHIKV genome and host stress response genes. Accordingly, host transcripts translationally upregulated during infection are enriched in suboptimal codons with low Codon Adaptation Index (CAI), a parameter that quantifies how well a gene codon usage aligns with the preferred codons of the genome under basal conditions. In contrast, translationally repressed genes are enriched in optimal codons with higher CAI values [8]. To explore whether similar mechanisms occur in mosquito cells, we analyzed the CAI of translationally activated and repressed transcripts, along with tRNA modification levels. Unlike in human cells, CHIKV infection in mosquito cells did not result in significant differences in CAI between groups or across time points in mosquito cells (**Fig 5**). This finding was further supported by the observation that the GC/AT content remains consistent across transcripts and at all codon positions (**S12 Fig**). We further validated these observations by examining tRNA modification levels at 8 hpi, when global translation is repressed but CHIKV RNA is translationally activated; 1 dpi, when the global translation begins to recover; and 7 dpi, representing the persistent infection stage. Our LC-MS/MS analysis of tRNA modifications revealed no significant changes in any of the examined modifications, including mcm⁵ (**S3 Table** and **S13 Fig**), at any of the infection time points. Collectively, these results indicate that CHIKV codon usage remains suboptimal in infected mosquito cells.

**Fig 5.**
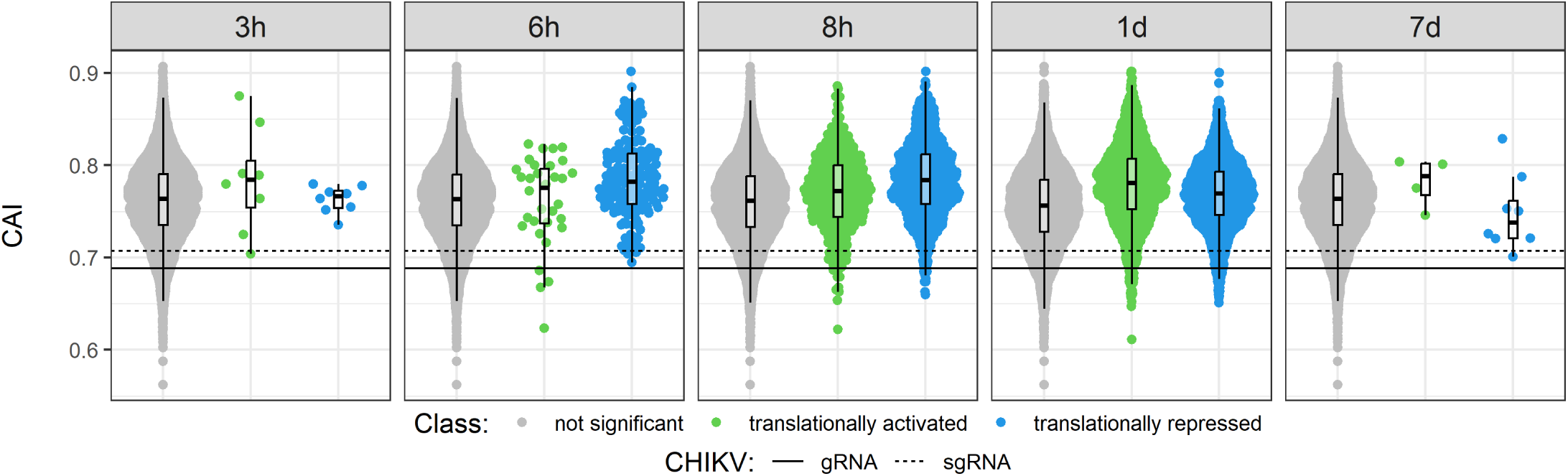
CHIKV infection does not alter host codon optimality. Representation of Codon adaptation index (CAI) at the different time-points post-infection. Each dot represents one gene. Significant translationally activated and translationally repressed genes are colored in green and blue, respectively. Horizontal lines indicate values for CHIKV ORFs: solid line for gRNA, dashed line for sgRNA.

## Discussion

The ability of arboviruses to persist in mosquito vectors without causing significant pathology is crucial for their transmission and widespread impact on global health. Our study, using CHIKV as a model in *A. albopictus* C6/36 cells, reveals a finely tuned translational virus-host equilibrium for sustained viral production through controlled viral protein synthesis. Moreover, this translational balance is independent of the RNAi pathway. A striking finding is the uncoupling of viral RNA and protein expression during persistent CHIKV infection in mosquito cells. Although viral RNA levels continue to rise or plateau, viral protein expression and progeny titers peak early and subsequently decline. Through orthogonal validation we demonstrate that this uncoupling results from a virus-specific translation repression, even at the peak of viral RNA accumulation (**Fig 1**, **Fig 2** and **Fig 4**). This indicates that, rather than maximizing viral output, CHIKV adopts a strategy of sustained but controlled viral production, ensuring long-term host cell viability. These findings stand in stark contrast to the highly efficient CHIKV protein synthesis observed in human cells (**Fig 6**) [8].

**Fig 6.**
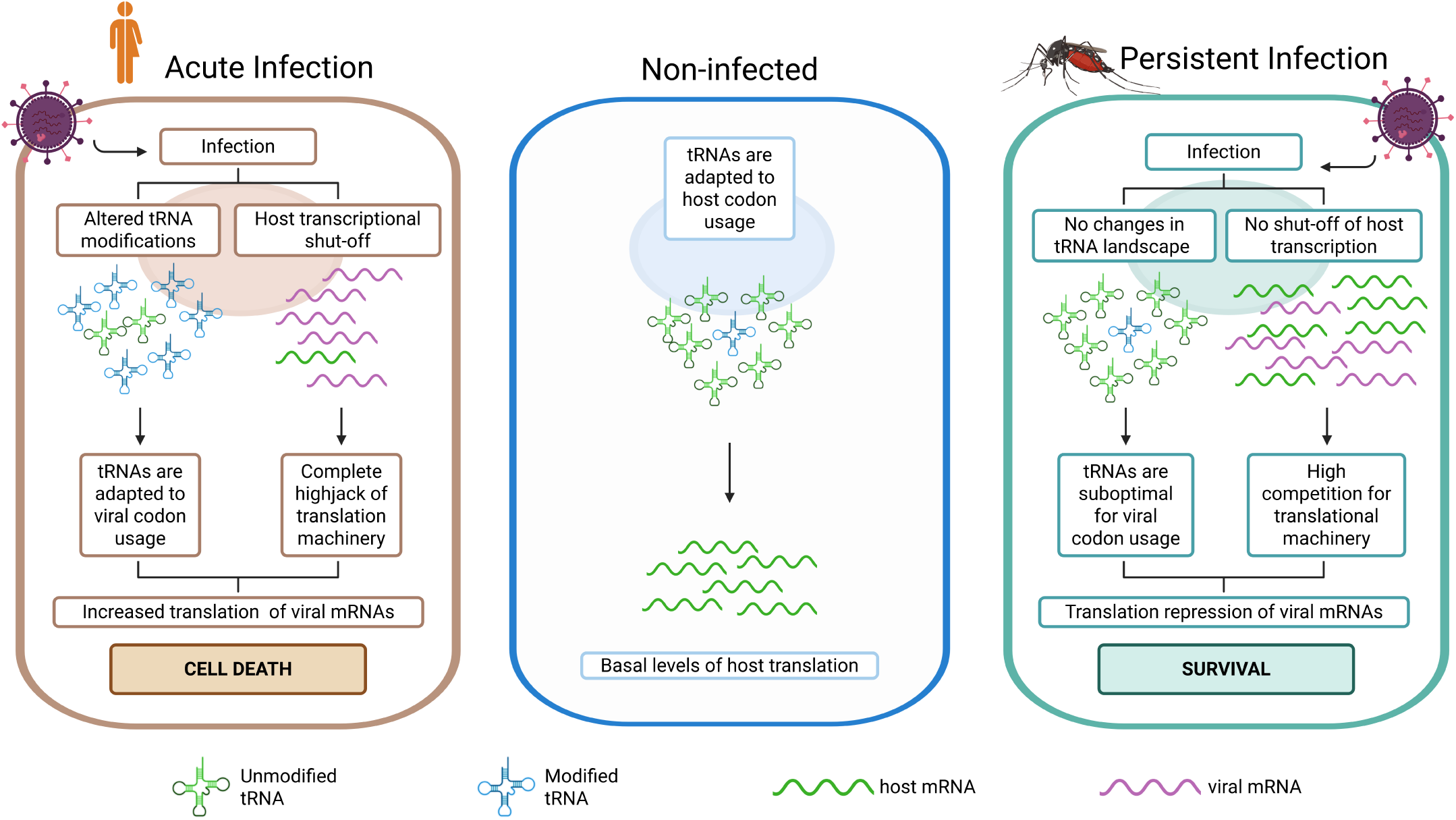
Model for CHIKV infection in humans and mosquitoes. In non-infected conditions of any organism, mRNA translation is adapted to preferentially express basal genes with optimal codons. These codons are efficiently translated as their corresponding tRNA concentrations are highly abundant. During acute infection of an arbovirus, as in human infection, replication triggers a stress response of the cell that will modify the tRNA anticodon. The changes in tRNA will favor the decoding of rare codons in which viral mRNAs are enriched. Hence, viral mRNAs are highly efficiently translated. In this situation, the viral infection will induce cellular death. In contrast, in mosquito cells the virus will establish a persistent infection. During persistence, the tRNA pool is not changed to favor viral codon usage, which will induce a moderate translation repression on viral mRNAs. This will allow the mosquito cell to survive the infection and maintain high levels of viral mRNAs.

Our study reveals two key distinctions in how the virus navigates critical determinants of translational efficiency in human versus mosquito cells: codon optimality and the competition for translational resources between viral and host mRNAs (**Fig 6**). First, CHIKV codon usage is inherently suboptimal for both human and mosquito hosts, predicting inefficient translation under baseline conditions. However, in human cells, CHIKV infection triggers a DNA damage response (DDR) that reshapes codon optimality, enhancing the translation of CHIKV RNAs, which are enriched in codons also found in DDR-associated genes [8]. As a result, CHIKV codon usage becomes effectively optimized under infection conditions. In contrast, in mosquito cells, comparisons of codon usage between translationally activated and repressed host mRNAs during CHIKV infection, along with tRNA modification analyses, revealed no changes in codon optimality, and CHIKV codon usage remains suboptimal (**S12-S13 Fig** and **Fig 5**). This observation aligns with the known high resistance of mosquito cells to DDR induction [16]. Second, CHIKV infection in human cells induces a dramatic reduction in host mRNA levels. This reduction is driven by the nuclear translocation of the viral NSP2 protein, which inhibits host mRNA transcription and makes limited translational resources, such as low abundant tRNAs, more available for viral translation. In mosquito cells, however, NSP2 remains cytoplasmic throughout infection, resulting in stable host mRNA levels. Together, these two features, persistent suboptimal codon usage and sustained host mRNA abundance, limit viral protein production in mosquito cells (**Fig 3** and **Fig 5**). We propose that these differences contribute critically to the distinct infection outcomes observed in humans and mosquitoes.

Based on our findings, we propose a refined model for CHIKV persistence in mosquito cells. During the early stages of infection (6 hpi to 1 dpi), CHIKV induces an acute host stress response characterized by a transient increase in eIF2α phosphorylation (**Fig 2D**). This phosphorylation causes a global, eIF2α-dependent attenuation of host mRNA translation, which subsequently triggers the activation of ATF4 and the transcriptional upregulation of ATF4-dependent genes associated with cell survival and metabolic adaptation. In contrast to host mRNAs, CHIKV RNA translation becomes progressively activated during this early phase. This active translation, particularly of sgRNA which is activated earlier than gRNA, indicates that CHIKV can evade eIF2α phosphorylation-mediated translational inhibition. This phenomenon is consistent with findings observed in other alphaviruses, such as Sindbis or Semliki Forest viruses, which employ structural features in their sgRNA to facilitate efficient translation initiation under conditions of eIF2α phosphorylation [17]. Moreover, the global repression of host translation liberates translational resources, including tRNAs, that are limited under physiological conditions, thereby further facilitating viral RNA translation. At the persistent stage, a dynamic shift occurs. Host mRNA levels and translation largely recovers, while CHIKV RNA translation is progressively downregulated. We propose that this repression in viral protein production is multifactorial, resulting from a return to an eIF2α-dependent translation initiation environment and increased competition for limited translational resources. This, combined with CHIKV suboptimal codon usage, which is not compensated by host tRNA modifications in mosquito cells, limits viral protein production to levels that supports sustained persistence without compromising host cell viability.

In summary, our study reveals a biphasic model of translational regulation underlying CHIKV persistence in mosquitoes, characterized by an early phase of active viral translation followed by a self-limiting repression that finely balances viral replication with host cell survival. These findings highlight the importance of translational control as a key determinant of persistent infection in mosquito vectors.

## Material and Methods

### Cell Lines and Virus

C6/36 (kindly provided by Andres Merits) and U4.4 (kindly provided by Emilie Pondeville) cells were grown at 28°C and cultivated in Leibovitz 15 medium (L15, ThermoFisher,11415064) supplemented with 10% Tryptose Phosphate Broth (TPB, BD, 260300) and 10% Fetal bovine serum (FBS, Merck, F7524-500ML). HEK293T (ATCC; CRL-11268) cells were grown at 37°C and 5% CO_2_ and cultivated in Dulbecco’s modified Eagle’s medium (DMEM, ThermoFisher Scientific, 41966029) supplemented with 10% FBS and 1% non-essential amino acids (ThermoFisher Scientific,11140035). CHIKV LR2006-OPY1 viral stock (GenBank: DQ443544, kindly provided by Andres Merits) was produced and titrated in HEK293T cells as in [8].

### Viral infections

All CHIKV infections of C6/36 and U4.4 cells were carried out with a MOI of 4, incubated for 1h at 28°C and harvested at the indicated time points after infection.

In C6/36 infections for RNA and protein analyses, 6-well plates were previously coated with poly-L-lysine (Merck, P1274-100MG) and then seeded with 5×10^5^ cells/well. On the next day, cells were washed once with PBS (Cytiva, SH30028.02) and then infected with CHIKV stock. After incubation, the virus was removed and 2 mL of pre-warmed media was added. At time of harvest, supernatant was taken and stored at –80°C until titration, cells were washed once with PBS and then lysed with 250 µl ice-cold lysis buffer (10 mM Tris-HCl pH 7.4, 10 mM MgCl_2_, 100 mM NaCl, 1 % Triton, 2 mM DTT). Cells were scraped and transferred to a new tube and snap-frozen with liquid nitrogen. For time points longer than 1 dpi, media was changed every 24 h; cells were jumped out in 1.5mL of fresh media at time of harvest and 1:5^th^ was re-plated in new poly-L-lysine pre-treated 6-well plates. The rest was centrifuged at 4°C 2500 rpm to pellet. The pellet was carefully washed with PBS, again centrifuged and finally lysed with 250 µl of ice-cold lysis buffer. U4.4 infections were carried out the same with the following differences: seeding for 6-well plates was of 3×10^5^ cells/well without poly-L-lysine coating and for time points longer than 1 dpi, cells were split 1:5^th^ every 48 h using trypsin 0.25% (ThermoFisher Scientific, 25200056) for 5 min.

For Ribosome and Polysome profiling analyses, T-150 plates previously coated with poly-L-lysine were seeded with 1×10^7^ cells. On the next day, cells were washed with PBS and then incubated with CHIKV. After 1h of incubation, the virus was removed and 15mL of pre-warmed media was added. At time of harvest, cells were previously incubated with non-supplemented media with 100 μg/mL cycloheximide (CHX, Merck, C7698), then washed once with PBS + 100 μg/mL CHX and finally lysed with 700ul ice-cold lysis buffer + 0.25% Deoxycholate and 100 μg/ml CHX. Cells were scraped and transferred to a new tube and snap-frozen with liquid nitrogen. For 5 and 7 dpi time points, cells were split 1:5^th^ into new poly-L-lysine pre-treated T-150 plates to achieve 80% confluency at time of harvest.

### RNA extraction, RT-qPCR, protein analyses and titer

For further processing, all samples were centrifuged at 4°C for 5 min at 12.000xg and lysates were transferred to new tubes for subsequent analysis.

Total RNA was isolated following proteinase K treatment (New England Biolabs, P8107S), phenol-chloropropane extraction, and ethanol precipitation. The pellet was resuspended in RNAse-free water and consecutively treated with 0,1µl Turbo DNAse/1μg RNA (ThermoScientific, AM1907) for 30 min at 37°C. 18 nanograms (ng) of total RNA was analyzed by TaqMan RT-qPCR using qScript XLT One-Step RT-qPCR ToughMix (Quanta BioSciences, 733-2232). The following probes and primers were used: CHIKV NSP1 Fwd (5’-aaccccgttcatgtacaatgc-3’), Rev (5’-gtacctgctcatctgcccaatt-3’) and probe (5’-6Fam-cgggtgcctacccctcatactcgac-TAMRA-3’); CHIKV Envelope Fwd (5’-aagctccgcgtcctttaccaag-3’), Rev (5’-ccaaattgtcctggtcttcct-3’) and probe (5’-6FAM-ccaatgtcttcagcctggacaccttt-TAMRA-3’). 1 µg of *in vitro* CHIKV-RNA was serially diluted 1:10 to generate a standard curve for absolute quantification of viral RNA molecules.

After BCA quantification of total lysate using Pierce BCA Protein Assay Kit (ThermoScientific, 23227), the equal volume of lysate corresponding to 25 µg was mixed with NuPAGE LDS Sample buffer (4x) (ThermoScientific, NP0007). Samples were then boiled for 5 min at 95°C and subsequently loaded onto a 10% polyacrylamide-SDS gel. Following electrophoresis, proteins were transferred to a membrane using TurboBlot (Biorad) for 7 min at 25V. Then the membranes were blocked with 5% non fat dry milk in TBST for 1 h at room temperature and then incubated overnight at 4°C with the primary antibodies. Antibodies used in western blotting were: CHIKV NSP1 (1:1500), Capsid (1:1500) and NSP2 (1:5000) (kindly provided by A. Merits), Tubulin (Merck, T6199-100UL), and P-eIF2α (1:1000, Cell Signaling, 9721S). Next day they were washed 3x for 5min with TBST and incubated at room temperature for 1 h with secondary antibodies (GE Healthcare, NA934V or NA931V) diluted in 5% milk in TBST. Subsequently, the membranes were washed again 3x for 5 min with TBST and exposed to ECL solution (Cytiva, GERPN2134D2 or ThermoFisher, 10005943). Band detection was done using the BioRad ChemiDoc MP Imaging System. Total band intensity volumes of proteins of interest were normalized to the housekeeping gene of the same sample and the mean of 3 biological replicates was calculated. Viral protein levels were further normalized to the highest intensity time point (1 dpi for C6/36 and 2 dpi for U4.4). P-eIF2α in infected samples was normalized to the mock levels of the protein at each corresponding time point.

Supernatants were titrated using standard plaque assay. Briefly, supernatants for each time point were diluted 1:10 in non-supplemented DMEM media. HEK293T cells were seeded at 3×10^5^ cells/w in 12-well plates and incubated with the diluted supernatants for 1 h at 37°C and shaken every 10 min. After incubation the virus was changed to pre-warmed DMEM + 2% FBS + 1% carboxymethyl cellulose (CMC, Merck, 21902) in a 1:1 ratio and put back into a 37°C incubator for 3 days. Then, cells were fixed and stained with crystal violet for 30 min under UV light. Finally, cells were rinsed with warm tap water and plaques were counted.

### Immunofluorescence

For immunofluorescence analysis, we seeded 2×10^5^ cells/well for C6/36 and 8×10^4^ cells/well for U4.4 into 24-well plates containing sterile glass coverslips coated with poly-L-Lysine. Cells were infected with CHIKV as explained before, and at different time-points post-infection, cells were processed for confocal study. Cells were washed with PBS and fixed with 4% paraformaldehyde (Sigma-Aldrich, P6148) for 15 min. Then, they were permeabilized with PBST (PBS + 0.5% Triton X-100 (Sigma-Aldrich X-100) for 20 min in ice. After a wash with PBS, cells were blocked in blocking buffer (PBS + 10% FBS) for 30 min at room temperature, washed, and then incubated 1 h with the different antibodies. Using blocking buffer, antibodies were diluted at: CHIKV proteins, kindly provided by Andres Merits, 1:10.000 for anti-Capsid (rabbit) and 1:1000 for anti-NSP2 (rabbit), and 1:2500 for anti-J2 (mouse) anti-double-stranded RNA (clone J2) (Nordic-MUbio, 10010200). Then, cells were washed twice with PBS and incubated with the secondary antibody anti-rabbit Alexa FluorTM 488 goat IgG and anti-mouse Alexa FluorTM 568 goat IgG (ThermoFisher, A-11004 and –11008) diluted 1:1000 in blocking buffer for 45 min covered from the light. Cells were washed and immersed in DAPI staining (1 µg/ml, Sigma-Aldrich) for 5 min, covered from the light. Lastly, slides were mounted using pre-warmed Mowiol (Sigma-Aldrich, 32459). Cells were imaged using a Leica SP5 inverted confocal laser scanning microscope and processed with Fiji (ImageJ). For percentage calculations, 4 randomly selected fields per slide were acquired from at least two biological replicates for each infection time point.

### Polysome Profiling

Sucrose (Sigma-Aldrich, 84097) solutions at 10% and 50% were prepared dissolved in polysome buffer 2.0 (20mM Tris-HCL pH 7.5, 10mM MgCl2, 100 mM NH4Cl), and supplemented with CHX 100ug/ml and DTT 2mM. 6mL of the 10% sucrose gradient was added to SW41Ti tubes (Beckman) and then 6mL of 50% sucrose was underlaid using a needle. To get a 15-45% linear gradient, we used the Gradient Master. Gradients were stored at 4°C overnight. Lysates were thawed on ice and centrifuged for 5 min at 12.000xg. 400 µL of lysate coming from the same number of cells, were loaded onto the linear sucrose gradients, and centrifuged for 3 h at 35.000rpm at 4°C in a Beckman SW41 rotor. BCA quantification was performed to normalize the amount of sample loaded into the gradient. Gradients were immediately fractionated into 20 fractions using a Biocomp Fractionator. Fractions of 500 µl were collected in 1,5mL RNAse-free tubes with 55 µL of 10% SDS.

Fractions were extracted using phenol-chloroform and precipitated with ethanol. RNA of fractions corresponding to 40S, 60S, 80S and polysome peaks were pooled together when dissolving with RNAse free water. Samples were DNAse treated before analyzing by qPCR. Viral RNA was assessed as described above using TaqMan qPCR.

### Ribosome Profiling libraries

Lysates for Ribosome Profiling were divided into two: half for mRNA-seq and half for Ribo-seq. 6 A260 units of lysate for Ribo-seq were digested with 250U of RNAse I (ThermoFisher Scientific, AM2294) for 5 min at 25°C. Digestion was stopped by adding inhibitor SUPERase-In (ThermoFisher Scientific, AM2694) double the volume of RNAse I. The digested sample was fractionated using a polysome profiling fractionator and the RNA of fractions corresponding to the monosome peak was extracted as described previously. The library was prepared following the protocol of Ingolia et al., 2012 [18]. Briefly, RNA of 25-35 nt was purified by gel, 3’ dephosphorylated with T4 PNK (NEB, M0201S), ligated to cloning linker (Biolabs, S1315S) and ribosomal RNA (rRNA) depleted with the Ribo-Zero Gold rRNA Removal kit (Human/Mouse/Rat, Epicenter/Illumina, MRZG126). Adapter-ligated-RNA was then reverse transcribed with Superscript III (ThermoFisher Scientific, 18080085) using the RT primer (5’-AGATCGGAAGAGCGTCGTGTAGGGAAAGAGTGTAGATCTCGGTGGTCGC-(SpC18)-CACTCA-(SpC18)-TTCAGACGTGTGCTCTTCCGATCTATTGATGGTGCCTACAG-3’) and again gel purified. CircLigase II (Epicenter, CL9025K) was used for circularization of the cDNA. Lastly, the library was generated with Phusion polymerase (NEB, M0530S) with PCR forward primer (5’-AATGATACGGCGACCACCGAGATCTACAC-3’) and Illumina index primers (NEBNext, E7335). The library was gel purified before sequencing. Lysates for mRNA-seq were RNA extracted with proteinase K (NEB, P8107S) and phenol-chloropropane extraction method and DNAse treated (ThermoScientific, AM1907). 2,5 µg of total RNA was used for library preparation with the NEBNext® Ultra™ II RNA Library Prep Kit for Illumina® (NEB, E7775S) and NEBNext Poly(A) mRNA Magnetic Isolation Module (NEB #E7490) following manufacturer’s instructions. Samples were sequenced with single-end 75bp reads for Ribo-seq and paired-end 75bp reads for mRNA-seq on an Illumina NextSeq500to reach at least 30 million reads per sample.

### Data analysis

Ribo-seq and RNA-seq adapter sequences were trimmed from raw reads using cutadapt (v1.18) [19], retaining reads with a minimum of 50 nt for RNA-seq and between 25-35 nt for Ribo-seq. Surviving reads were mapped to an index of known tRNA, rRNA, snRNA, snoRNA and miRNA genes based on the non-coding sequences for *A. albopictus* txid7160 downloaded from the European Nucleotide Archive (ENA-EBI) via bowtie (v1.2.3) [20], and only non-mapping reads were kept. Since eukaryotic transcripts usually feature several different splice isoforms that can rise to different ORFs, we determined the most prominent mRNA isoform for each gene present in our RNA-seq samples to allow a gene-based analysis. We aligned the RNA-seq reads using kallisto (v0.43.0) [21] against *A. albopictus* genome and transcriptome (*Aedes albopictus Foshan*, AaloF1 rel.48, www. vectorbase.org) [22] and quantified abundance with sleuth (v0.29.0) [23]. The protein coding isoform which on average was expressed highest was selected. Subsequently, CHIKV genome and transcriptome (LR2006 OPY1 GenBank KT449801.1) was combined with the mosquito reference. mRNA and RPF reads were aligned to the final *A. albopictus* annotation using STAR (v2.7.0) [24]. Quality control for RPF libraries were assessed as the following. Read lengths of 29-33 nt were selected which contained the full range of ribosome footprints (**S4 Fig**). For each of the read lengths in 29-33 nt, the offset of the A site from the 5’ end of reads was calibrated using the start codons of CDS [25], which indicated the following A-site offsets: 16 nt offset for 29-31 nt read lengths, and 17 nt offset for 32 nt read lengths (i.e. 13 and 14 nt offset for P sites, respectively). For metagene plots in **S6-10 Fig**, the ribosome density of P sites at each position relative to the start or stop codon was divided by the overall density along the CDS; the average across all CDSs with more than 50 reads was taken. For **S11 Fig**, for each of the read lengths in 29-32 nt, we computed the ribosome counts of P sites in each of the three possible translation frames along the CDSs (excluding the first and the last 15 nt).

Counts for RNA-seq reads were quantified using featureCounts (subread v1.5.1) [26], while assignment to A– and P-site and CDS footprint counting was performed using riboWaltz (v2.0) [27]. Read length of 29-33 nt was selected and 14–18 nt were used as corresponding P-site offsets across all samples. Before continuing our analysis, all genes with fewer than a total of 5 RPF and 10 RNA-seq reads were discarded. In addition, since read mapping to the CHIKV sgRNA region can originate from both the full-length CHIKV gRNA (NSP ORF) as well as the sgRNA (SP ORF), we calculated the average coverage across the NSP specific region to estimate how many read counts mapping to the SP region should originate from the full-length (=NSP) isoform, assuming a uniform sequencing coverage. Using these estimates, we then re-assigned the number of reads from the SP to the NSP ORF for each sample prior to the differential expression analysis. Once RNA-seq and Ribo-seq reads had been quantified, we directly compared the corresponding counts for each sample to ensure linear correlation between RPF and mRNA counts (**S5 Fig**). Lastly, we assessed replicate quality by principal component analysis for the RNA-seq and Ribo-seq counts (**S3 Fig**).

For differential analyses of host expression signatures, the counts of viral genes were excluded. Differential expression analysis of RNA-seq and Ribo-seq counts was performed using DESeq2 [28]. Translation efficiency was computed with RiboDiff [14]. GO enrichment analysis was performed using the “enrichr” function of the clusterProfiler package [29], against the GO annotations from the VectorBase database.

We used an orthology-based approach to functionally characterize *A. albopictus* genes with unknown functions. First, we ran Broccoli (v1.2) [30] to build gene orthologies of protein-coding genes between *A. albopictus* and (i) the set of twenty bilaterian species and (ii) the set of eight insect species considered in [31]. For *A. albopictus*, we used the GTF from AaloF1 (rel.48, www.vectorbase.org), while for all other species we used the GTFs from [31]. In order to avoid redundant gene orthology calls, we selected one representative protein isoform for each gene in each species (i.e., the isoform with the longest protein-coding sequence). From the orthologous_pairs.txt file returned by Broccoli for each run, we selected all orthologous pairs involving *A. albopictus* and (i) *Homo sapiens / Mus musculus* genes in the bilaterian set or (ii) *Aedes aegypti / Drosophila melanogaster* from the insect set. Finally, based on these orthologies correspondences, we assigned the annotated gene description of the *H. sapiens/ M. musculus* (annotations from NCBI assembly GRCh38.p14 and GRCm39) or *A. aegypti / D. melanogaster* (annotations from www.vectorbase.org AaegL5.3 LVP_AGWG, rel. 68, and NCBI assembly Release 6 plus ISO1 MT, respectively) genes to the corresponding *A. albopictus* orthologs. The generated functional annotation and the relative sources are reported in **S1 Table**.

Codon adaptation indexes for each of the transcripts were calculated via the CAI function from the seqinR package (v4.2-36) [32] using weights based on all genes in the counts matrix. Frequencies of GC nucleotides per codon position was calculated by first finding the occurrence per codon in each transcript using seqinR function uco (index “eff”) and then calculating the sum of those codons with the same nucleotide for position and divided by total number of codons. Percentage of GC content was calculated as the sum of nucleotide occurrence and divided by total length of the cds for each transcript. Results were visualized using ggplot2.

For comparison with human infection, we used publicly available ribosome profiling data of CHIKV infection in HEK293T [8]. Raw counts for mRNA and RPFs were downloaded from NCBI Gene Expression Omnibus (GEO) (GSE143390). Counts for cytoplasm and endoplasmic reticulum fraction coming from the same sample were added before calculating the proportion of genes in each dataset.

### Quantification of tRNA modifications by LC-MS/MS

3-10 μg of total RNA per sample was run in a 15% TBE-UREA gel (Novex, ThermoFisher Scientific, EC6885BOX) for 60 min at 180V. Gels were stained with 1:10000 SybrGold (ThermoFisher Scientific, S11494) and the tRNA bands were selected. tRNA was then extracted using ZR small RNA PAGE Recovery kit (Zymo Research, R1070) following the manufacturer’s instructions. The same amount of extracted tRNA was digested with 1 μl of the Nucleoside Digestion Mix (New England BioLabs, M0649S) and the mixture was incubated at 37°C for 1 h. Samples (15 ng) were analyzed using an Orbitrap Eclipse Tribrid mass spectrometer (Thermo Fisher Scientific, San Jose, CA, USA) coupled to an EASY-nLC 1200 (Thermo Fisher Scientific (Proxeon), Odense, Denmark). Ribonucleosides were loaded directly onto the analytical column and were separated by reversed-phase chromatography using a 50 cm column with an inner diameter of 75 μm, packed with 2 μm C18 particles (Thermo Fisher Scientific, ES903). Chromatographic gradients started at 93% buffer A and 3% buffer B with a flow rate of 250 nL/min for 5 min and gradually increased to 30% buffer B and 70% buffer A in 20 min. After each analysis, the column was washed for 10min with 0% buffer A and 100% buffer B. Buffer A: 0.1% formic acid in water. Buffer B: 0.1% formic acid in 80% acetonitrile. The mass spectrometer was operated in positive ionization mode with nanospray voltage set at 2.4kV and source temperature at 305°C. For Parallel Reaction Monitoring (PRM) method the quadrupole isolation window was set to 1.4m/z, and MSMS scans were collected over a mass range of m/z 50-300, with detection in the Orbitrap at resolution of 60,000. MSMS fragmentation of defined masses (**S3 Table**) was performed using HCD at NCE 20 (except stated differently, **S3 Table**) [33], the auto gain control (AGC) was set to “Standard” and a maximum injection time of 118 ms was used. In each PRM cycle, one full MS scan at resolution of 120,000 was acquired over a mass range of m/z 220-700 with detection in the Orbitrap mass analyzer. Auto gain control (AGC) was set to 10^5^ and the maximum injection time was set to 50 ms. Serial dilutions were prepared using commercial pure ribonucleosides (0.005-150 pg, Carbosynth, Toronto Research Chemicals) in order to establish the linear range of quantification and the limit of detection of each compound. A mix of commercial ribonucleosides was injected before and after each batch of samples to assess instrument stability and to be used as an external standard to calibrate the retention time of each ribonucleoside.

Acquired data were analyzed with the Skyline-daily software (v24.1.1.284) and extracted precursor areas of the ribonucleosides were used for quantification.

## Data availability

Source data are provided with this paper for Ribo-seq and RNA-seq have been submitted to the NCBI Sequence Read Archive (SRA) under accession number SUB15400962. The raw proteomics data have been deposited to the EMBL-EBI MetaboLights database with the identifier REQ20250616211235.

## Acknowledgements

This work has received funding by “la Caixa” Foundation (LCF/PR/HR23/52430003), the Spanish Ministry of Science and Innovation (PID2022-136939OBI00/MICIN/AEI/10.13039/501100011033 and “ERDF a way of making Europe”), by the 2021 SGR 00176 grant from the Departament de Recerca i Universitats de la Generalitat de Catalunya and by an institutional “Unidad de Excelencia María de Maeztu,” funded by the MCIN and the AEI (CEX2024-001431M). We acknowledge support of the Spanish Ministry of Science and Innovation through the Centro de Excelencia Severo Ochoa (CEX2020-001049-S, MCIN/AEI /10.13039/501100011033), and the Generalitat de Catalunya through the CERCA programme. MP-T was a recipient of an FPI fellowship from the Spanish Ministry of Science and Innovation (PRE2020-093049). The work of X.H. has been supported by a PhD fellowship from the Fundación Ramón Areces. We are grateful to the UPF and CRG Core Technologies Programmes for their support and assistance in this work. We thank R. Böttcher and R. Barrios for insightful discussions and E. Muscolino and A. Meyerhans for critically reading the manuscript and for helpful discussions. Figures 1A and 6 were created using BioRender for which we own a full licence.

## Supporting Figures/Tables Captions

**S1 Fig.**
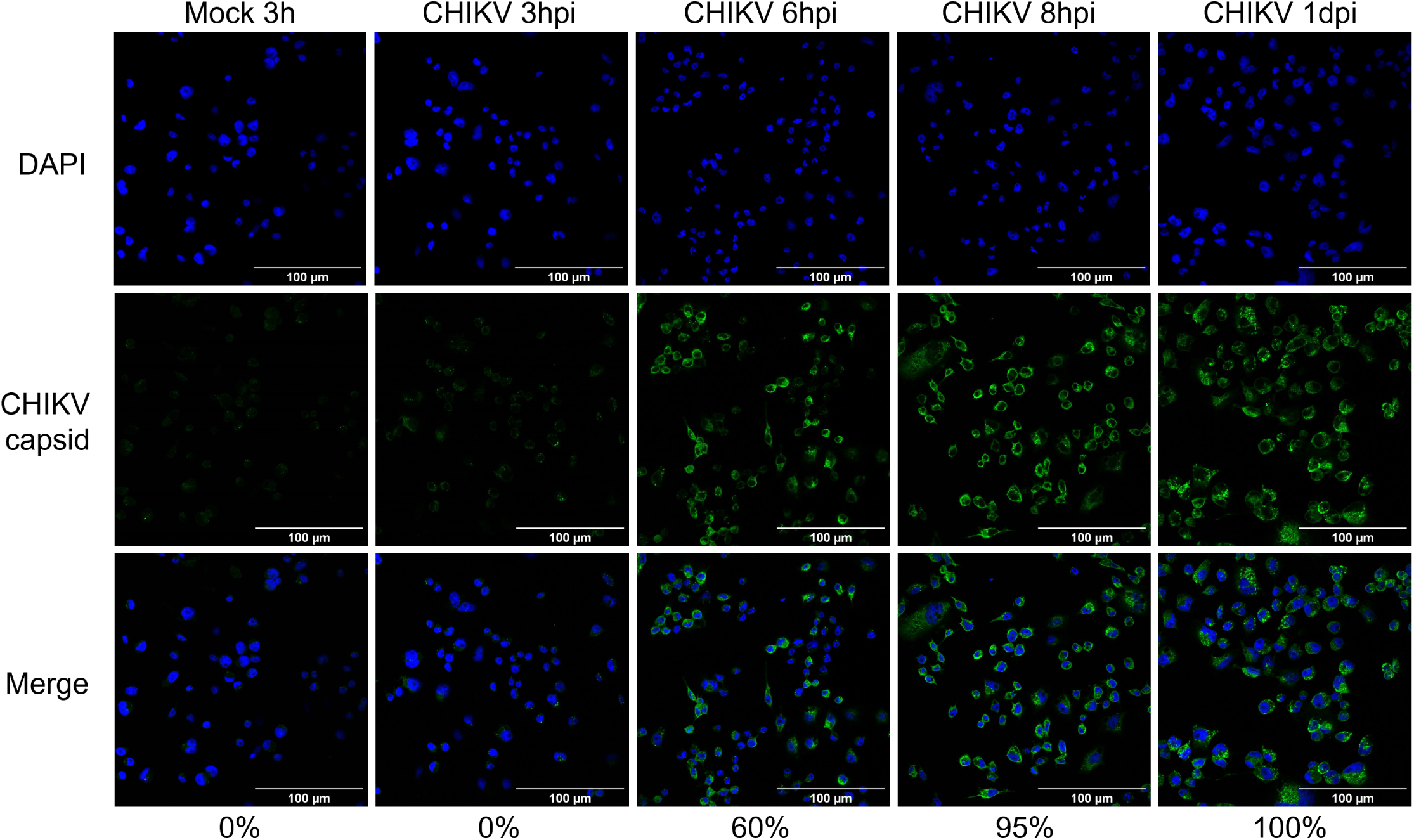
Percentage of CHIKV-infected C6/36 cells during the course of infection. Immunofluorescence staining was used to monitor CHIKV infection in C6/36 cells. Infected cells were labeled with anti-CHIKV capsid antibody (green), and nuclei were stained with DAPI (blue). Scale bars correspond to 100 μm. Percentage indicates the proportion of cells positive for capsid CHIKV-protein.

**S2 Fig.**
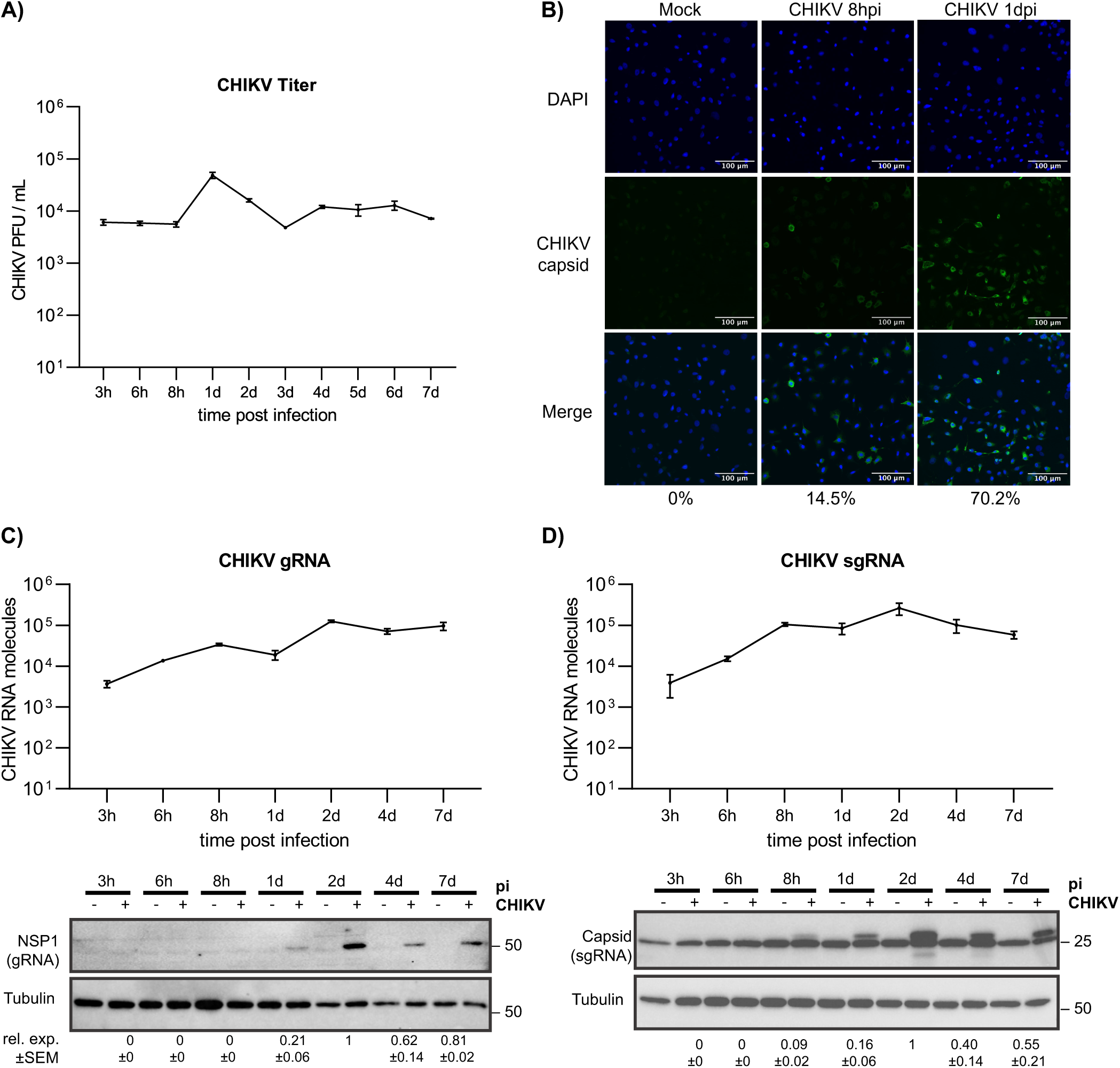
Kinetics of CHIKV infection in U4.4 mosquito cells. (A) CHIKV titers over time, determined by plaque assay. (B) Immunofluorescence staining was used to monitor CHIKV infection in U4.4 cells. Infected cells were labeled with anti-CHIKV capsid antibody (green), and nuclei were stained with DAPI (blue). Scale bars correspond to 100 μm. Percentage indicates the proportion of cells positive for capsid CHIKV-protein. (C) Infection kinetic of CHIKV gRNA levels determined by qPCR and NSP1 protein levels assessed by western blotting. Tubulin was used as loading control. (D) Infection kinetic of CHIKV sgRNA levels and capsid protein expression, analyzed as in panel (C). All infections were performed at a MOI of 4 and samples were collected at the indicated time post-infection. Data points represent the mean ± SEM of 3 independent biological replicates. Viral RNA levels were quantified using a standard curve of an *in vitro* generated CHIKV RNA. Protein expression levels correspond to the intensity quantification of each band normalized to tubulin levels and to the 2 dpi value within each blot.

**S3 Fig.**
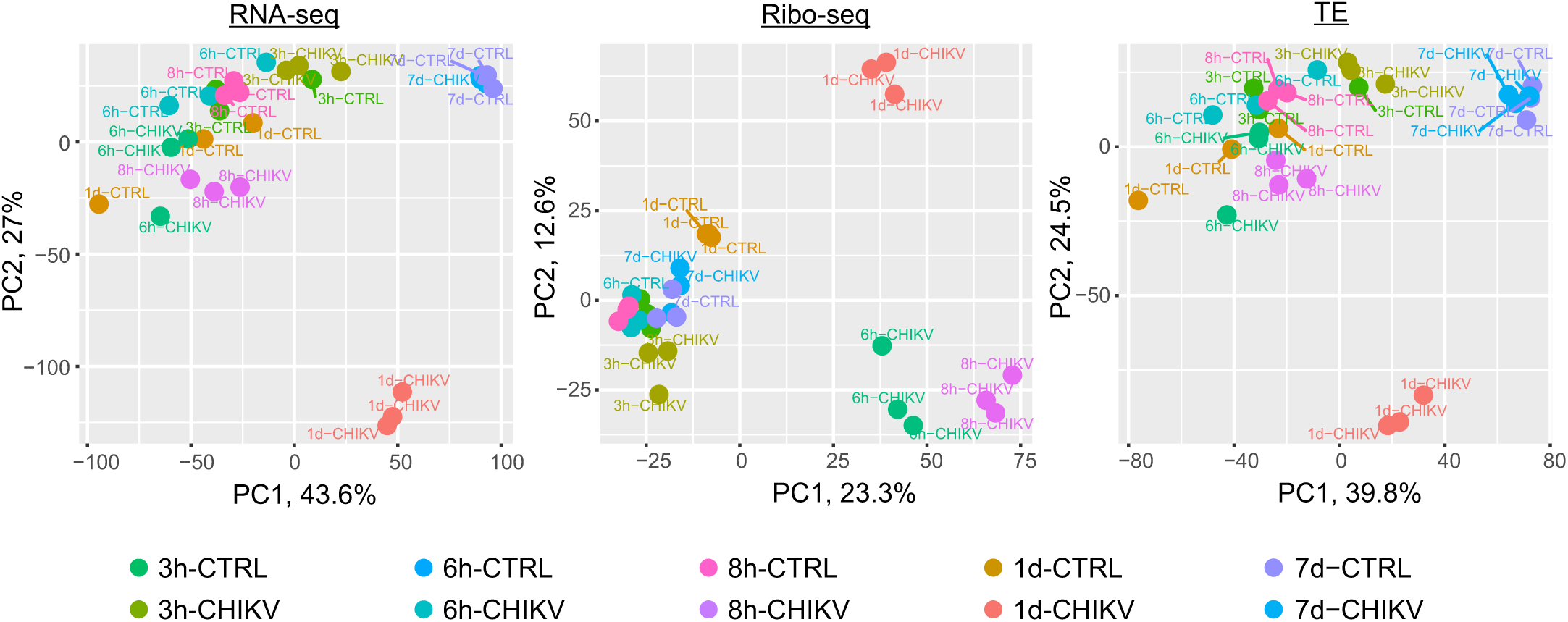
Principal component analysis (PCA) of RNA-seq, Ribo-seq, and translational efficiency (TE) data. PCA of the log-10 transformed counts from RNA-seq and Ribo-seq datasets, as well as on calculated TE values, to assess sample clustering and variability across conditions.

**S4 Fig.**
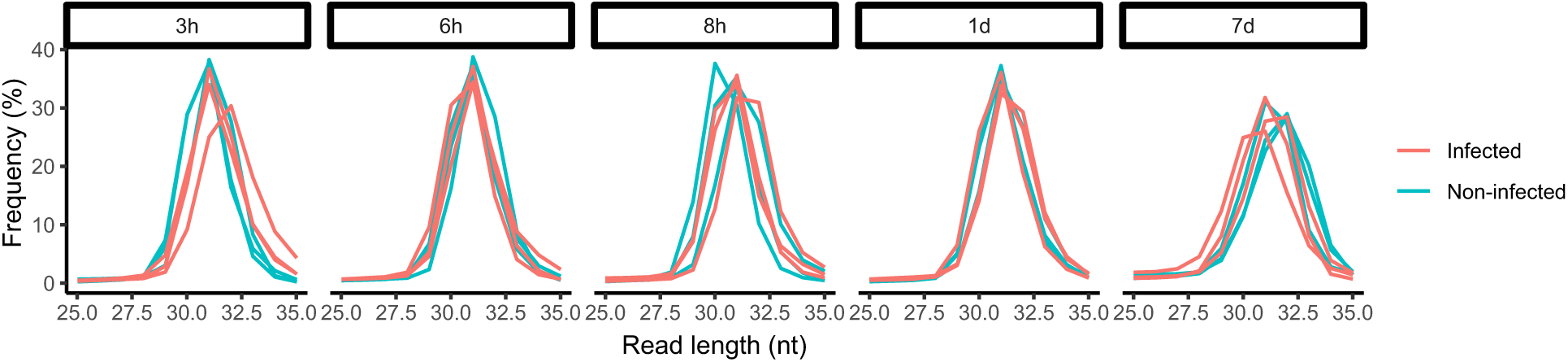
Distribution of ribosome-protected fragment (RPF) lengths. Length distribution of RPFs at each time post-infection in CHIKV-infected and non-infected conditions. Each replicate is represented separately.

**S5 Fig.**
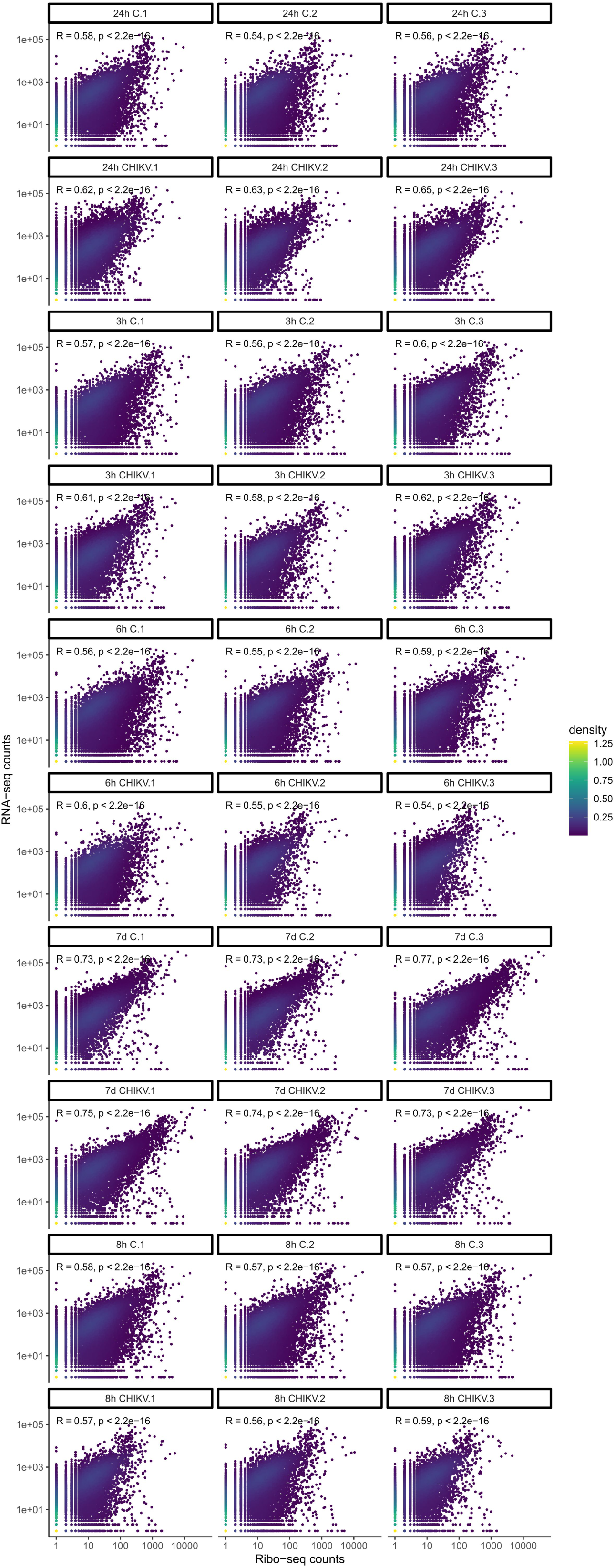
Comparison of RNA-seq and Ribo-seq libraries. Scatter plots depicting the correlation between RNA-seq and Ribo-seq counts for each gene across all samples. The color represents the point density.

**S6 Fig.**
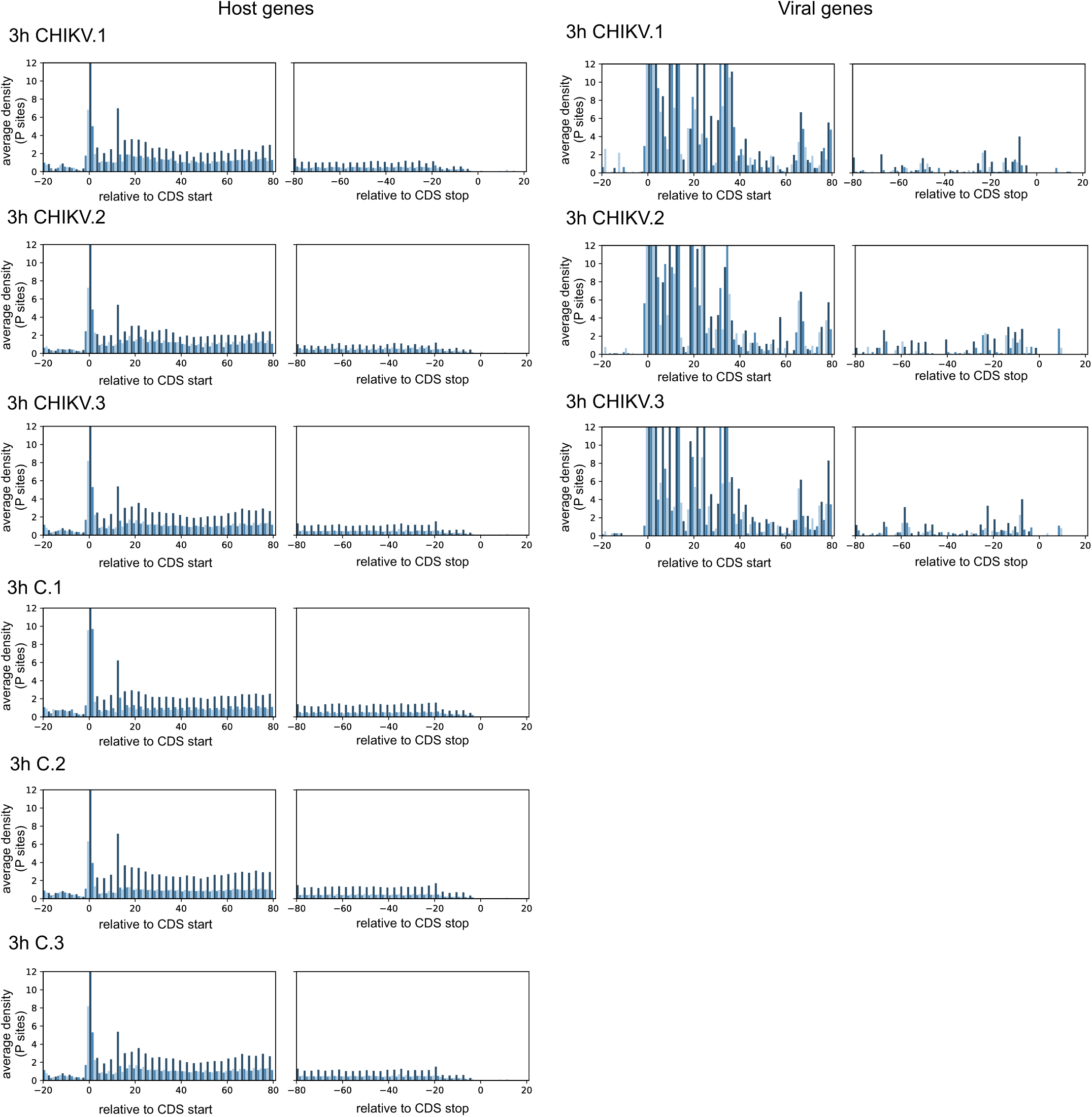
Metagene distribution in 3 hpi samples. RPFs triplet periodicity at 5’ UTR and 3’ UTR for (A) host and (B) viral derived RPFs.

**S7 Fig.**
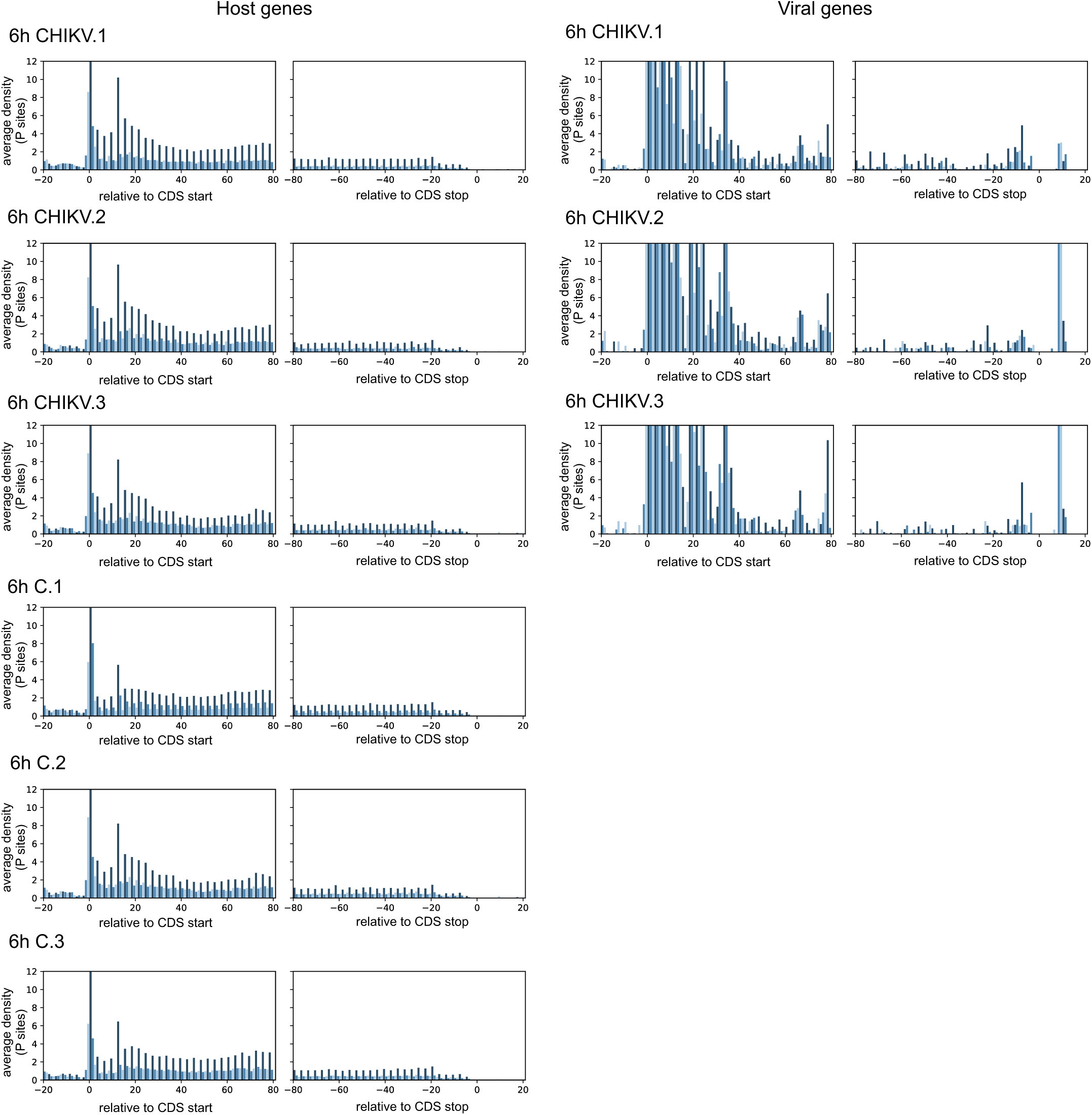
Metagene distribution in 6 hpi samples. RPFs triplet periodicity at 5’ UTR and 3’ UTR for (A) host and (B) viral derived RPFs.

**S8 Fig.**
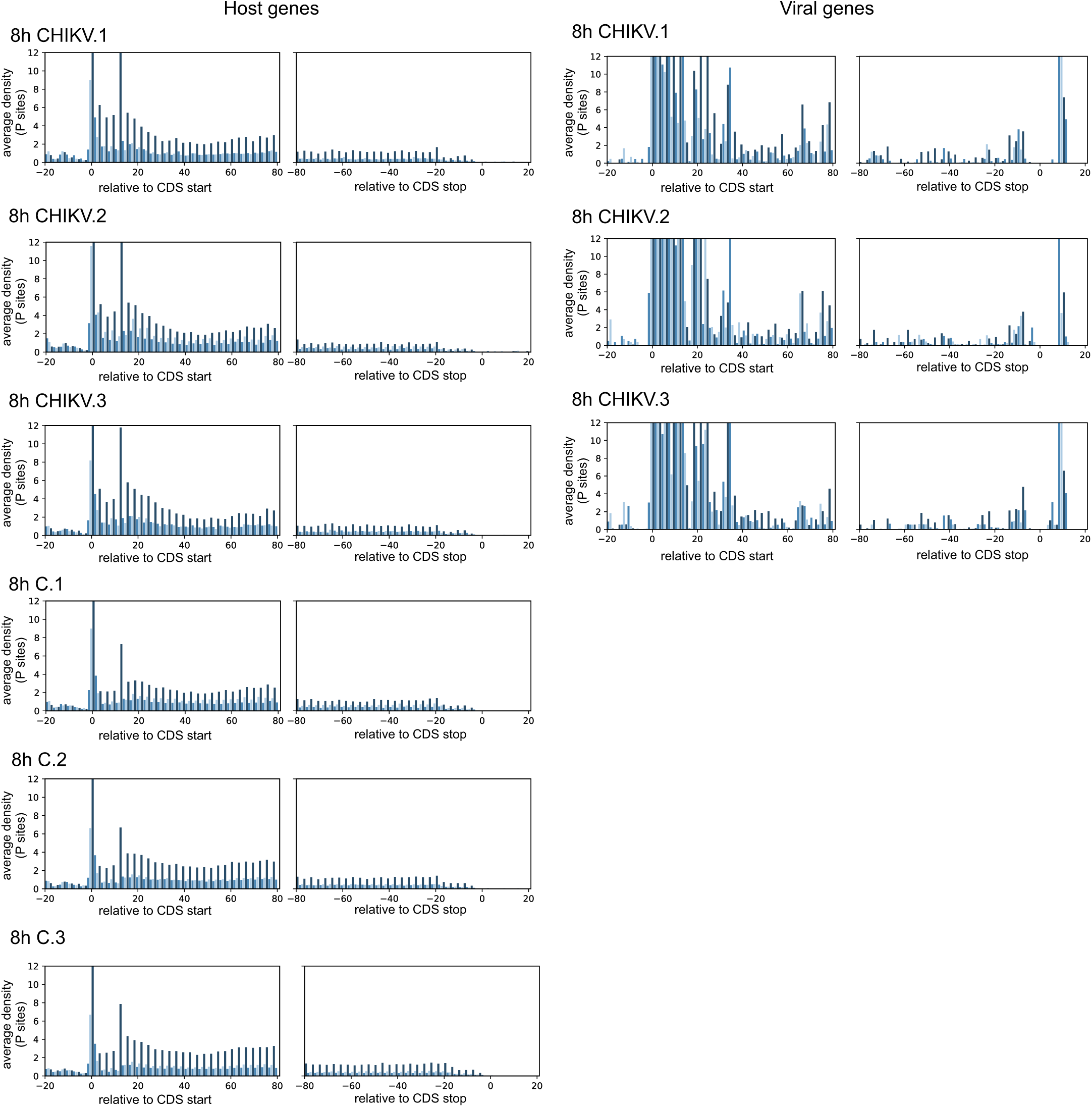
Metagene distribution in 8 hpi samples. RPFs triplet periodicity at 5’ UTR and 3’ UTR for (A) host and (B) viral derived RPFs.

**S9 Fig.**
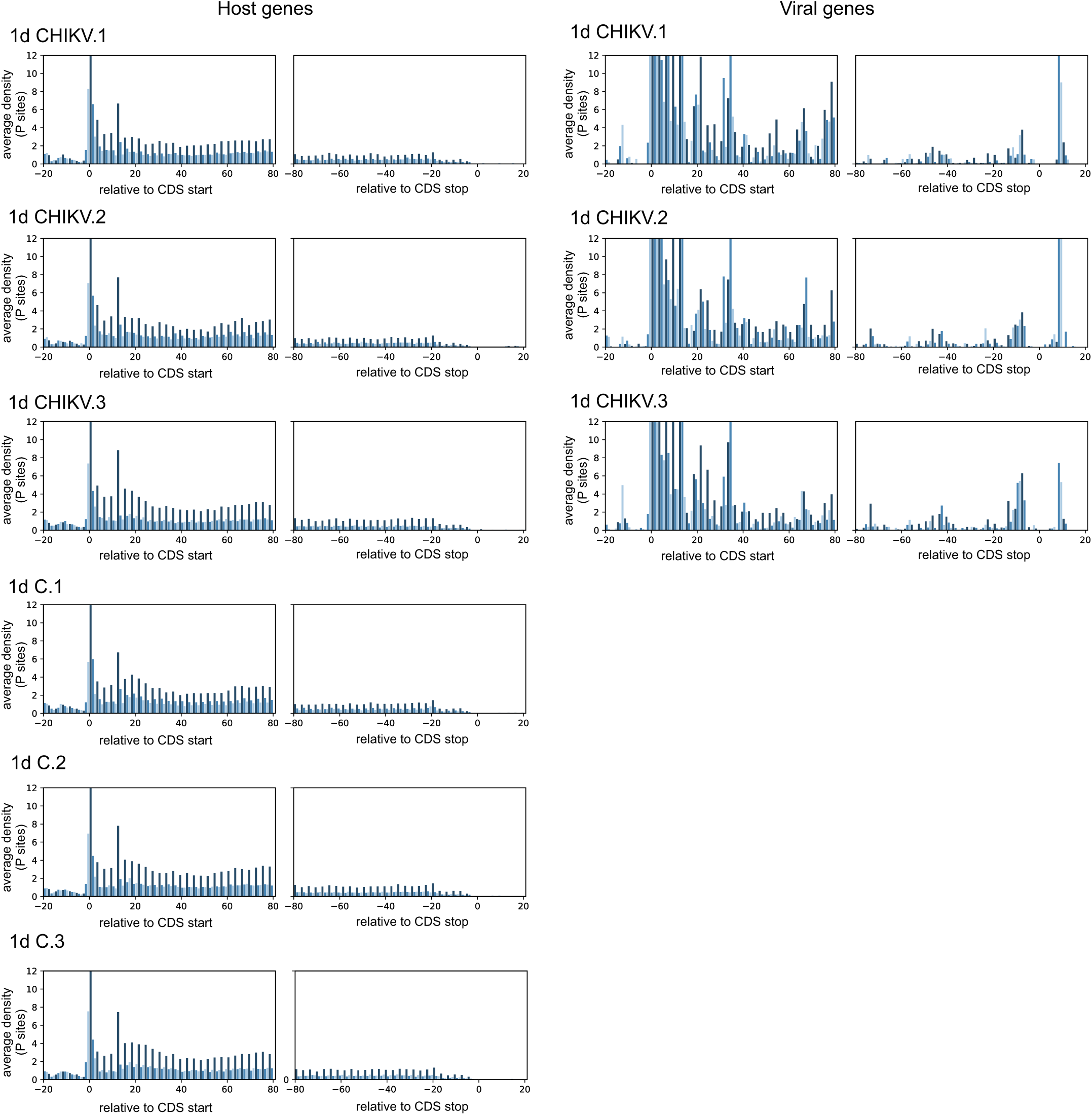
Metagene distribution in 1 dpi samples. RPFs triplet periodicity at 5’ UTR and 3’ UTR for (A) host and (B) viral derived RPFs.

**S10 Fig.**
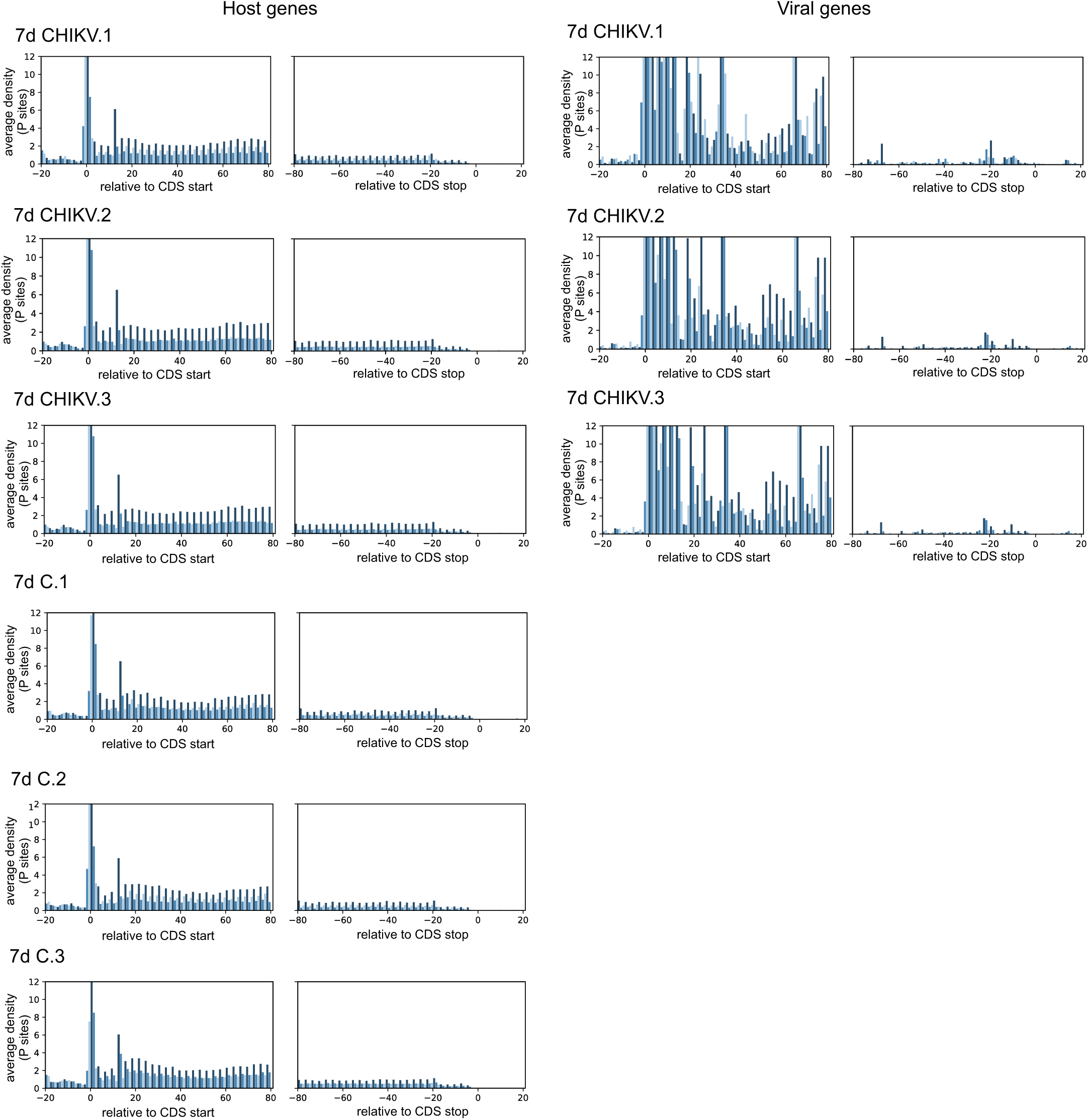
Metagene distribution in 7 dpi samples. RPFs triplet periodicity at 5’ UTR and 3’ UTR for (A) host and (B) viral derived RPFs.

**S11 Fig.**
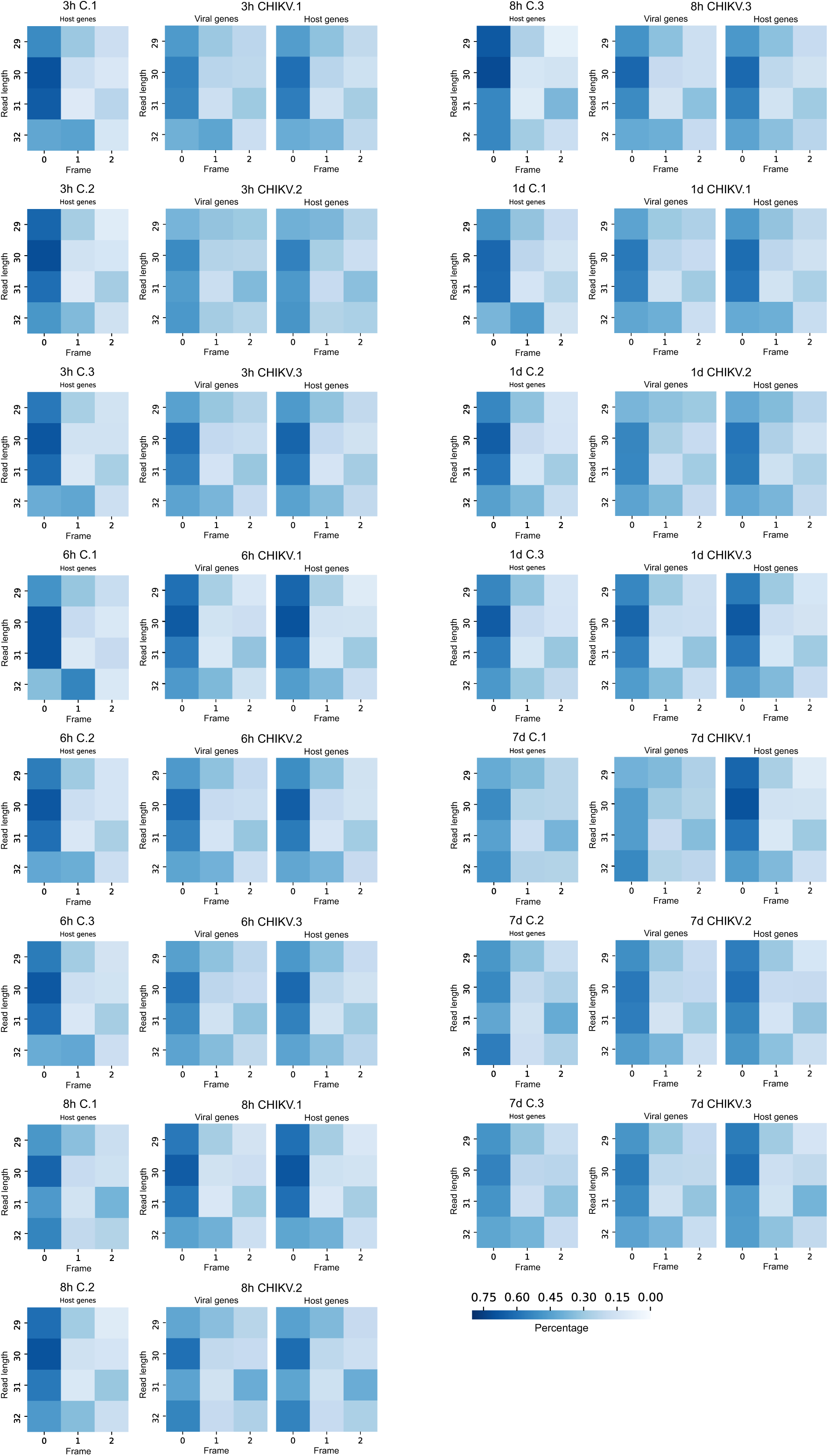
P-site frame distribution in Ribo-seq samples. Distribution of RPF p-site across the three possible reading frames. Color represents percentage of reads

**S1 Table.** Comprehensive analyses of differentially expressed genes for RNA-seq and translation efficiency. Columns represent: gene ID –as per vectorbase nomenclature–, gene product description, log2 fold change (log2FC) and p-adjusted value (padj) of CHIKV-infected versus non-infected mRNA-seq counts, mRNA class –whether transcription of the RNA was upregulated, downregulated or not significantly altered in CHIKV infection-, log2FC and padj of CHIKV-infected versus non-infected RPF-seq counts, gene translation efficiency (TE) in mock samples, gene TE in infected samples, log2FC and padj of CHIKV-infected versus non-infected TE, TE class –whether translation efficiency was activated, repressed or not significantly altered in CHIKV infection-, and ortholog product description based on insect (drosophila) or vertebrate (*Homo sapiens* and *Mus musculus*) bilateral analyses as described in Materials and Methods.

**S2 Table.** GO terms for classes of differentially expressed genes at the level of mRNA-seq and TE. Each library is stored in separate sheets. GO terms were analyzed for the following gene expression groups: UP (mRNA upregulated), DOWN (mRNA downregulated), TA (TE translationally activated) and TR (TE translationally repressed). Columns represent gene expression group (Cluster), GO term number (ID), GO term description (Description), ratio of genes in the cluster associated with the GO term (GeneRatio), background ratio (BgRatio), p-value, adjusted p-value, false discovery rate (qvalue), list of associated gene IDs –as per vectorbase nomenclature– (geneID), and the number of genes associated with the GO term (Count).

**S12 Fig.**
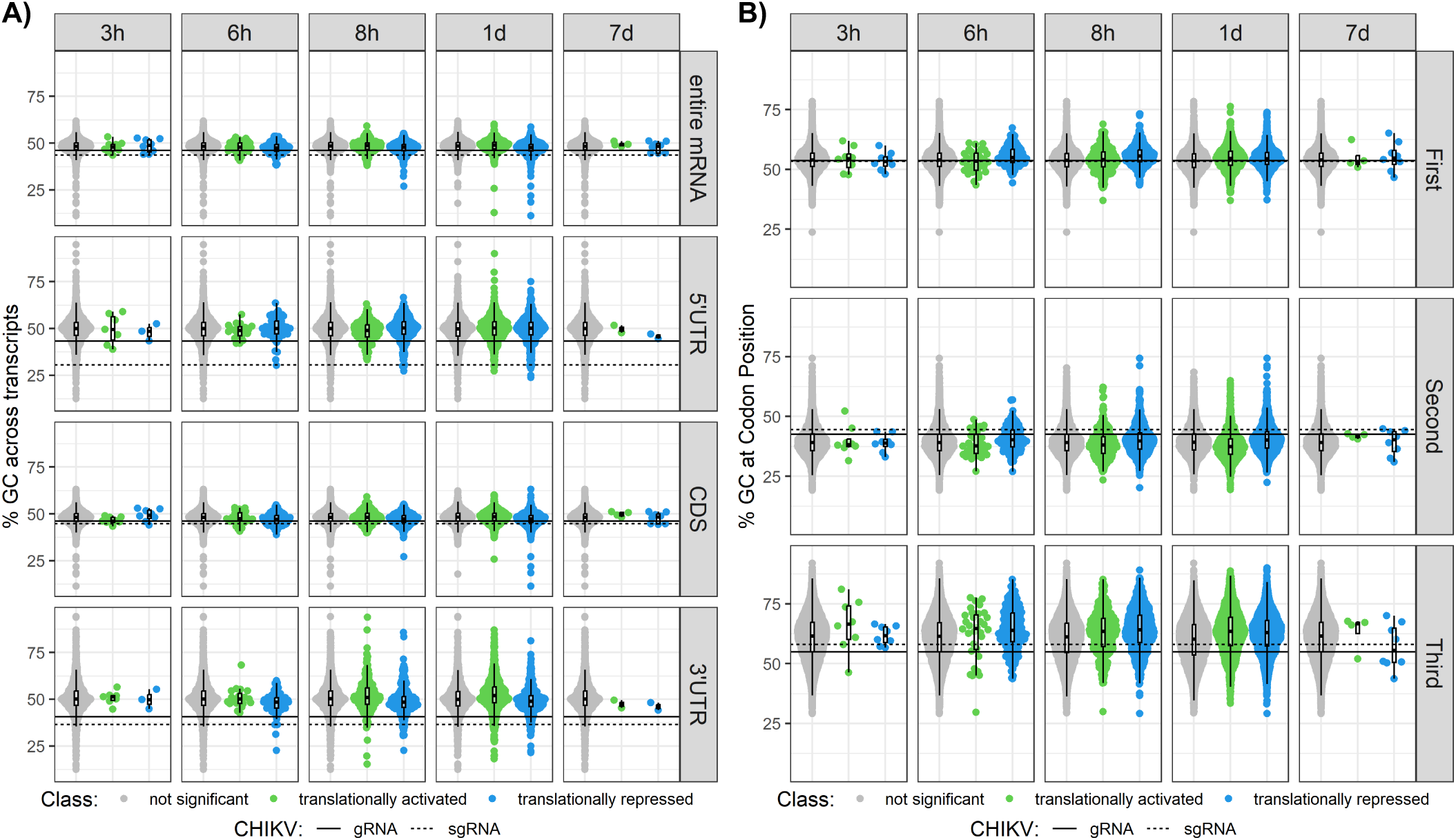
CHIKV infection does not reprogram codon optimality. (A) GC content distribution in transcripts at the 3’ UTR, 5’ UTR, CDS or entire mRNA sequence among different translational groups across time points. (B) GC content at codon position (first, second or third position) among different translational groups across time points. Each dot represents one gene. Significant translationally activated (TA) and translationally repressed (TR) genes are colored in green and blue, respectively. Horizontal lines indicate CHIKV ORF values: solid line for gRNA, dashed line for sgRNA.

**S3 Table.** List of tRNA modifications. List of all modified ribonucleosides identified in mass spectrometry analyses of tRNA samples with defined masses of its MSMS fragmentation.

**S13 Fig.**
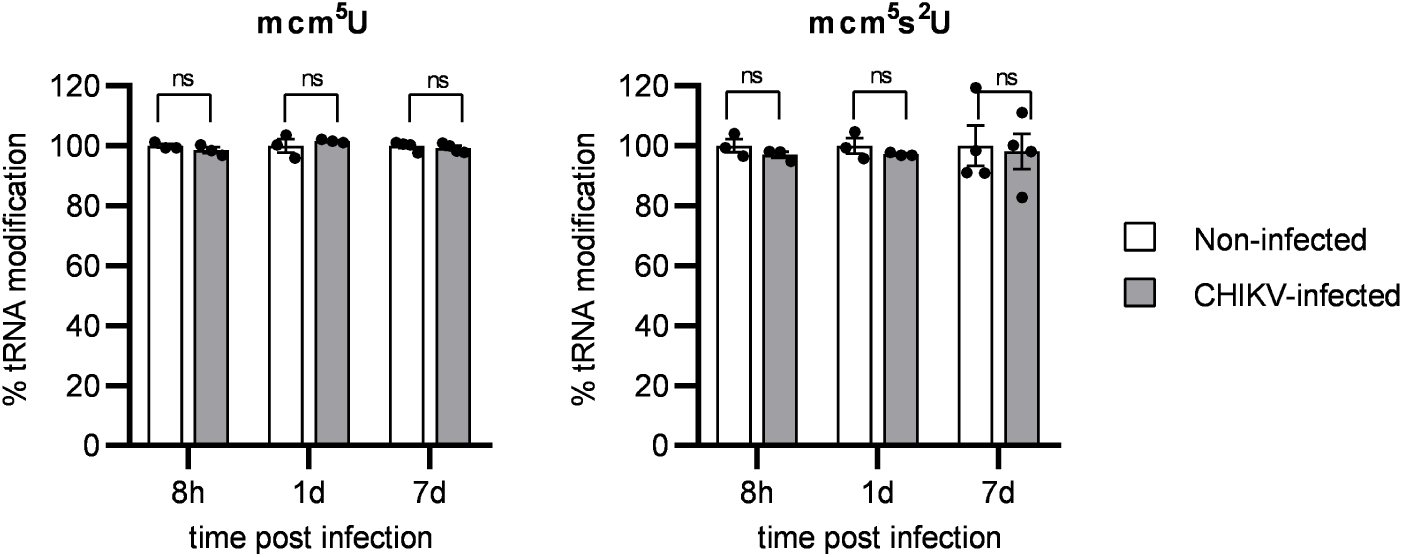
CHIKV infection does not reprogram the tRNA epitranscriptome. Relative levels of mcm^5^U and mcm^5^s^2^U tRNA modifications in CHIKV-infected cells compared to non-infected cells, measured by LC-MS/MS. Bars represent the mean ± SEM of 3 independent biological replicates.

## Notes

### Competing Interest Statement

The authors have declared no competing interest.

